# Patterns of Interploidy Admixture in Polyploid Complexes: Insights from *Thymus* sect. *Mastichina* (Lamiaceae)

**DOI:** 10.1101/2025.11.27.690785

**Authors:** Francisco José García-Cárdenas, María Ángeles Ortiz, José Carlos del Valle, David Doblas-Pruvost, Manuel de la Estrella, Diego Nieto-Lugilde, Lisa Pokorny, Regina Berjano

**Affiliations:** Departamento de Biología Vegetal y Ecología, Universidad de Sevilla, Avda. Reina Mercedes s/n, 41012 Seville, Spain; Departamento de Botánica, Ecología y Fisiología Vegetal, Universidad de Córdoba, Campus de Rabanales, Edificio Celestino Mutis, E-14071 Córdoba, Spain; Real Jardín Botánico (RJB), CSIC, 28014 Madrid, Spain

**Author notes:** Lisa Pokorny and Regina Berjano should be considered joint senior authors. **Correspondence**: Regina Berjano.

**Keywords:** Angiosperms353, Hyb-Seq, Interploidy Admixture, Phylogenomics, Polyploid Bridges, Population Genomics

## Abstract

Understanding gene flow between ploidy levels in polyploid complexes is essential for species delimitation and conservation. This study explores evolutionary dynamics in the polyploid complex *Thymus* sect. *Mastichina* (Lamiaceae), comprising three taxa: *T. mastichina* subsp. *mastichina*, *T. mastichina* subsp. *donyanae*, and the endangered *T. albicans*. Using Hyb-Seq data, phylogenomics (nuclear orthologs), and population genomics (SNPs), we confirm the section consists of two sister groups with distinct ploidy levels: a diploid and a tetraploid one. The tetraploid group shows low genetic differentiation among its populations, probably indicating rapid expansion across diverse environments. In contrast, the diploid group exhibits more complex genetic structuring, potentially shaped by geomorphology, interploidy introgression, and incipient isolation. Four diploid subgroups (Algarve, Cádiz, Doñana, and Hercynian) are identified, with reticulate evolution. The dense reticulation observed is compatible with incomplete lineage sorting in diploid lineages, due to recent and rapid divergence events. Phylogeographic analyses suggest isolation-by-distance, with two major riverbeds maybe playing a role in shaping genetic differentiation, while interploidy gene flow detected could have facilitated ancient and/or ongoing admixture between diploid and tetraploid lineages, despite geographic isolation. These findings highlight cryptic genetic diversity and emphasise the need for an integrative taxonomy that includes multiple lines of evidence: morphological, cytological, genomic, and ecological. Conservation efforts should prioritise protecting the four diploid subgroups, aided by flow cytometry, since they may harbour critical adaptive potential to both specific habitat types and/or environmental conditions. This work contributes to advancing our knowledge of evolution in polyploid complexes, by combining genomic approaches and highlighting cryptic diversity in *Thymus* species. Future research should investigate morphometric and chemical data, hybridisation events, divergence times, diversification dynamics, and relationships with other *Thymus* species to further understand polyploid evolution and its impact on biodiversity.

## 1 Introduction

Understanding the mechanisms driving speciation is a central focus of evolutionary biology. Polyploidy is considered one such mechanism of speciation with profound implications for plant evolution and ecology, which is mediated through whole-genome duplication (WGD; Van de Peer *et al*. 2017, 2021). Although this process has often been regarded as an evolutionary ‘dead end’, due to the limited evidence of ancient polyploidisation events (Arrigo and Barker 2012; Mayrose *et al*. 2015; but see Leebens-Mack *et al*. 2019), the existence of many polyploids of relatively recent origin supports its potential role as a driver of plant diversification (Vanneste *et al*. 2014a, 2014b; Soltis *et al*. 2015; Van de Peer *et al*. 2017, 2021). Even considering that polyploid lineages frequently face extinction, it seems that at least in rare cases they successfully establish and undergo significant diversification, as the “rarely successful hypothesis” postulates (Arrigo and Barker 2012; Hagen and Beaulieu 2024).

Polyploidy can arise through WGD within a single species (autopolyploidy) or after hybridisation (allopolyploidy), a well-established and prevalent process in plants that facilitates gene flow between distinct lineages that contributes to the emergence of novel species (Soltis and Soltis 2009; Sheidai and Koohdar 2023). Although newly established polyploids may initially experience reduced fertility and fitness due to genomic instability or changes in gene expression and epigenetic changes (Comai 2005; Porturas *et al*. 2019), once these issues are overcome, their potential for rapid adaptation can confer a selective advantage. At short-term, the success of polyploids can be explained by increased genetic variation, mediated by changes in gene expression and epigenetic remodelling, which influence the morphology, physiology, and ecology of newly formed polyploids (Schoenfelder and Fox 2015), and also by increased genetic robustness (Van de Peer *et al*. 2017). These changes, in turn, may affect interspecies interactions, confer tolerance to a broader range of ecological and environmental conditions, facilitate the colonisation of new habitats, and increase the potential for invasiveness (te Beest *et al*. 2012; Soltis *et al*. 2014; Van de Peer *et al*. 2021).

Polyploidy can promote sympatric speciation in plants, by acting as a natural reproductive barrier, as offspring from newly formed polyploids and their lower-ploidy progenitor(s) often exhibit unbalanced chromosome numbers, leading to sterility (the so-called *triploid block*; Ramsey and Schemske 1998; Otto and Whitton 2000; Adams and Wendel 2005). Nevertheless, this barrier to interploidy hybridization, it is sometimes leaky (for a review see Salony *et al*. 2025). Thus, the establishment of polyploid species in sympatry likely depends on prezygotic barriers between newly formed polyploids and their diploid progenitor(s), along with shifts from cross- to self-pollination or from sexual to asexual reproduction, which can mitigate cytotype minority exclusion of recently formed polyploids (Van de Peer *et al*. 2017). However, reproductive barriers following WGD may not be complete, particularly in cases where intermediate cytotypes are viable, which could facilitate the backcrossing with parental cytotypes (the so-called *triploid bridge pathway*; Ramsey and Schemske 1998; Husband 2004; Bartolić *et al*. 2025). Such backcrossing events may enable bidirectional gene flow, if the formed fertile interploidy hybrids are capable of backcrossing with their parentals (Kolář *et al*. 2017; Karbstein *et al*. 2022; Hodač *et al*. 2023; Bartolić *et al*. 2024). Otherwise, unreduced gametes could also mediate unidirectional gene flow from diploids to tetraploids, thus enabling interploidy gene flow (Leal *et al*. 2024).

Interploidy gene flow between lineages has turned out to characterise the evolution of many polyploid complexes (Nieto Feliner *et al*. 2020 and references therein), in which species with different ploidy levels interchange genetic material without apparently altering the chromosome number. Introgression and polyploidisation, although generally addressed in the literature as separate biological phenomena, are not mutually exclusive and often coexist in nature (Leal *et al*. 2024). These introgression events allow receptor species to accumulate new alleles not only through introgression with closely diploid progenitors, but also from more distantly related diploid or polyploid donor lineages, contributing to genetic diversity and facilitating adaptive allele emergence (Suarez-Gonzalez *et al*. 2018; Seear *et al*. 2020; Nieto Feliner *et al*. 2020; Schmickl and Yant 2021; Nibau *et al*. 2022). Despite growing recognition of their interplay, significant gaps remain in our understanding of how interploidy admixture in polyploid complexes shape evolutionary trajectories. Additionally, as polyploid complexes and introgression often blur phylogenetic relationships and obscure phenotypic distinctions (along with other factors such as incomplete lineage sorting, ILS), traditional taxonomic species differentiation can be challenging (Mallet 2007; Soltis *et al*. 2009; Al-Shehbaz 2012; Han *et al*. 2015). This is especially relevant for cryptic and threatened species because accurate identification assures the correct assessment and implementation of conservation strategies (Bickford *et al*. 2007; Fišer *et al*. 2018; Hending 2025).

Polyploidisation, introgression, and hybridisation, are common phenomena in several plant families, including Lamiaceae, and particularly in thyme—genus *Thymus* L. (e.g., Šoštarić *et al*. 2012; Yousefi *et al*. 2015). This genus is notable for its high species diversity (WFO 2025), significant economic value, high level of endemisms, and ecological importance, as thyme formations represent one of the most characteristic communities of Mediterranean shrublands (Morales 1986, 2002, 2010). In the Iberian Peninsula, *Thymus* sect. *Mastichina* presents an excellent model for studying evolutionary patterns driven by polyploidisation and introgression processes, which could be shaping relationships among their components. The section includes two closely related species (three taxa), whose populations are sometimes quite challenging to differentiate morphologically: *Thymus mastichina* (L.) L. and *T. albicans* Hoffmanns. & Link. *Thymus mastichina* is one of the most widely distributed endemic thyme species in the Iberian Peninsula. This species is known to hybridise with a large number of taxa with overlapping ranges (Morales 1986, 2010). Two subspecies have been described within it: *T. mastichina* subsp. *mastichina* (tetraploid), an Iberian widespread taxon, and *T. mastichina* subsp. *donyanae* (diploid), whose populations are mainly located within Doñana National and Natural Parks, in SW Spain (Morales 2010). The other species is *T. albicans* (diploid), found along the southwestern coast of the Iberian Peninsula across habitats highly threatened by coastal urban development (Valdés Castrillón *et al*. 1999). Taxonomists have differentiated these three taxa using morphological traits, mainly based on inflorescence diameter and calyx length (Morales 1986, 2002). However, this group exhibits a high degree of phenotypic variation, at both morphological (Morales 2010; Méndez-Tovar *et al*. 2015) and chemotypic levels (e.g., Miguel *et al*. 2003; Filipe *et al*. 2017). Therefore, unravelling the intricate relationships of this polyploid complex—essential for clarifying its taxonomic status—should be a priority, as it would help guide future conservation strategies (Hending 2025).

As High-Throughput Sequencing (HTS) technologies have become more accessible (Metzker 2010), population genomics methods have become an essential tool for reconstructing the evolutionary patterns that shape complex species histories (DePristo *et al*. 2011; Ellegren 2014). Among HTS techniques, genome skimming and target-enrichment have revolutionised our understanding of plant systematics (Mamanova *et al*. 2010; Dodsworth 2015; Pezzini *et al*. 2023). Hyb-Seq, which combines these two approaches (Weitemier *et al*. 2014; Dodsworth *et al*. 2019), has gained prominence in plant phylogenomics due to its affordability (reasonable per-sample cost), reproducibility, and replicability, while offering high resolution regardless of tissue-template quality (Villaverde *et al*. 2018; Brewer *et al*. 2019; Hale *et al*. 2020; Shee *et al*. 2020; Zuntini *et al*. 2024). While other genomic techniques, such as restriction site-associated DNA sequencing (RADseq) or genotyping-by-sequencing (GBS), have been used extensively to explore population structure and genetic diversity (Baird *et al*. 2008; Elshire *et al*. 2011; Peterson *et al*. 2012; Campos *et al*. 2024), the potential application of Hyb-Seq in population genetics still remains largely underexplored (Baker *et al*. 2021; McDonnell *et al*. 2021; but see Ousmael and Hansen 2024, Woudstra *et al*. 2025, or Viruel *et al*. 2025 for recent applications). However, current computational tools in phylogenomics and population genomics have been shown to effectively bridge micro- and macroevolutionary levels, using the same Hyb-Seq data, which can be encoded as either nuclear ortholog sequences or single-nucleotide polymorphism (SNPs; Beck *et al*. 2021; Slimp *et al*. 2021; Wenzell *et al*. 2021; Cooper *et al*. 2023; Phang *et al*. 2023; Tiley *et al*. 2024; Rincón Barrado *et al*. 2024a, 2024b; Balant *et al*. 2025; Pardo Otero *et al*. 2024; Freund *et al*. 2025; Liang *et al*. 2025; Stefani *et al*. 2025). By combining these analytical methods, insights into genetic connectivity, introgression, and the microevolutionary patterns of populations can be gained, which can be useful for identifying populations at risk of inbreeding or loss of adaptive potential and help in conservation and biodiversity management.

In this study, we aim to understand the evolutionary dynamics and genetic structure underlying the *Thymus* sect. *Mastichina* polyploid complex, with particular focus on the extent and patterns of interploidy admixture. To achieve this, we employ Hyb-Seq data in combination with phylogenomics and population genomics approaches to (i) resolve phylogenetic relationships among taxa in this section, (ii) assess genetic differentiation and admixture among diploid and tetraploid taxa within the complex, (iii) investigate signals of gene flow and introgression within and across ploidy levels, including potential pathways leading to interploidy admixture, and (iv) evaluate the implications of these patterns for taxonomy and conservation of *Thymus* sect. *Mastichina* taxa. This integrative framework seeks to clarify processes at the micro-macroevolutionary frontier shaping species boundaries in the presence of gene flow across ploidy levels in megadiverse Mediterranean scrubland endemics.

## 2 Materials and Methods

### 2.1 Study Group and Taxon Sampling

*Thymus* sect. *Mastichina* comprises two species and three taxa, all endemic to the Iberian Peninsula, differing in their distribution patterns across the region. *Thymus mastichina* subsp. *mastichina*, is a tetraploid taxon (2n = 56, 58, 60) widely distributed throughout the Iberian Peninsula across a variety of habitats such as roadside slopes, abandoned farmlands, pine forests, and mountain scree, growing on loose siliceous substrates but also on calcareous soils. *Thymus mastichina* subsp. *donyanae* is a diploid taxon (2n = 30) mainly located in coastal sandbanks and stabilised dunes within Doñana National and Natural Parks, in SW Spain (Huelva province). *Thymus albicans* is also a diploid taxon (2n = 28, 30) inhabiting stabilised subcostal dunes and coastal pine forests on sandy substrates along the coastline in Cádiz province (Spain) and Algarve (Portugal), SW Iberian Peninsula (Morales 2010). *Thymus albicans* is included in the Spanish national and regional Red Lists (Valdés Castrillón *et al*. 1999; Cabezudo *et al*. 2005), and is as well listed in the International Union for Conservation of Nature (IUCN) Red List (Buira *et al*. 2017).

We sampled a total of 40 populations from May to June 2023 across the Iberian Peninsula (Table 1). Twenty of those populations were the same studied in Girón *et al*. (2012). Individuals of each population were identified following Flora iberica (Morales 2010). Herbarium vouchers of each population were deposited at SEV herbarium (Table 1). Leaves of 8-12 individuals per population were dried in silica gel and stored at room temperature prior to DNA extraction. Additionally, we included three other species of *Thymus* from MA herbarium vouchers as outgroups (*T. carnosus* Boiss., *T. capitellatus* Hoffmanns. & Link and *T. camphoratus* Hoffmanns. & Link).

**TABLE 1.** Voucher information for studied populations, including identification according to Flora Iberica (currently recognised morphospecies), location, geographic coordinates, DNA content, ploidy-level estimation, and herbarium voucher numbers. Asterisks (*) indicate phenotypic variability in calyx length, both among and within individuals, which hinders reliable identification based on this character alone (*sensu* Morales *et al*. 2010).

| Species | Population ID | Population abbreviation | Locality | Lat | Long | C DNA (pg) | Ploidy | Herbarium voucher # |
| --- | --- | --- | --- | --- | --- | --- | --- | --- |
| *Thymus albicans* | A01 | PUERTO REAL | Spain. Cádiz, Puerto Real. Dehesa de las Yeguas. | 36.556 | -6.135 | 1.59 ± 0.01 (8) | 2x | SEV290208 |
| *Thymus albicans* | A02 | LA BARROSA | Spain. Cádiz, la Barrosa Suburban Park. | 36.363 | -6.170 | 1.60 ± 0.04 (9) | 2x | SEV290209 |
| *Thymus albicans* | A03 | ROCHE | Spain. Cádiz, Conil de la Frontera. ZEC Pinar de Roche. | 36.316 | -6.133 | 1.63 ± 0.02 (10) | 2x | SEV290210 |
| *Thymus albicans* * | A04, D03 | LA JUNCOSILLA | Spain. Sevilla, Villamanrique. La Juncosilla. Doñana Natural Park. | 37.192 | -6.329 | 1.54 ± 0.02 (15) | 2x | SEV290225 |
| *Thymus albicans* | A05 | FARO | Portugal. Algarve, Faro. Ria Formosa Natural Park. | 37.028 | -7.977 | 1.60 ± 0.02 (9) | 2x | SEV290212 |
| *Thymus albicans* | A06 | VILLARRASA | Spain. Huelva, Villarrasa. Dehesa del Duque. | 37.356 | -6.604 | 1.50 ± 0.01 (5) | 2x | SEV290213 |
| *Thymus albicans* | A07 | BONARES | Spain. Huelva, Bonares. | 37.259 | -6.725 | 1.52 ± 0.01 (10) | 2x | SEV290214 |
| *Thymus albicans* * | A08 | PINAR ALGAIDA | Spain. Doñana Natural Park. Pinar de la Algaída. | 36.855 | -6.308 | 1.53 ± 0.01 (11) | 2x | SEV290215 |
| *Thymus albicans* | A09 | QUARTEIRA | Portugal. Algarve, Quarteira. Vale do Lobo. | 37.062 | -8.080 | 1.54 ± 0.03 (8) | 2x | SEV290216 |
| *Thymus albicans* | A10 | VALE DO GARRAO | Portugal. Algarve, Almancil. Vale do Garrao. Ria Formosa Natural Park. | 37.044 | -8.038 | 1.55 ± 0.04 (7) | 2x | SEV290217 |
| *Thymus albicans* | A11 | NIEBLA | Spain. Huelva, Niebla. Rabo Conejo. | 37.489 | -6.719 | 1.55 ± 0.02 (11) | 2x | SEV290218 |
| *Thymus albicans* | A12 | LAS LOMAS | Spain. Cádiz, Vejer de la Frontera. Las Lomas. | 36.299 | -5.909 | 1.54 ± 0.05 (8) | 2x | SEV290219 |
| *Thymus albicans* | A13 | EL SOTO | Spain. Cádiz, Vejer de la Frontera. El Soto. | 36.242 | -5.929 | 1.6 ± 0.02 (9) | 2x | SEV290220 |
| *Thymus albicans* | A14 | CHICLANA | Spain. Cádiz, Chiclana. Pinar del Hierro. | 36.388 | -6.122 | 1.57 ± 0.02 (8) | 2x | SEV290221 |
| *Thymus albicans* * | A15 | VILLAMANRIQUE | Spain. Sevilla. Villamanrique de la Condesa. Doñana Natural Park. Dehesa de Gato. | 37.192 | -6.329 | 1.61 ± 0.03 (6) | 2x | SEV290222 |
| *Thymus mastichina subsp. donyanae* * | D01 | ALMONTE | Spain. Huelva, Almonte. El Porretal. ZEC Doñana North and West. | 37.167 | -6.497 | 1.50 ± 0.02 (9) | 2x | SEV290223 |
| *Thymus mastichina subsp. donyanae* * | D02 | HINOJOS | Spain. Huelva, Hinojos. ZEC Doñana North and West. | 37.287 | -6.446 | 1.51 ± 0.01 (10) | 2x | SEV290224 |
| *Thymus mastichina subsp. donyanae* * | D04 | PINHEROS DO MARIM | Portugal. Algarve, Quelfes. Pinheiros do Marim. Ria Formosa Natural Park. | 37.034 | -7.818 | 1.44 ± 0.02 (9) | 2x | SEV290226 |
| *Thymus mastichina subsp. donyanae* * | D05 | ABALARIO | Spain. Huelva, Abalario. Doñana Natural Park. | 37.072 | -6.651 | 1.51 ± 0.02 (8) | 2x | SEV290227 |
| *Thymus mastichina subsp. donyanae* * | D06 | RIBETEHILLOS | Spain. Huelva, Ribetehilos. Doñana Natural Park. | 37.150 | -6.643 | 1.51 ± 0.01 (10) | 2x | SEV290228 |
| *Thymus mastichina subsp. donyanae* * | D07 | CORRAL LIEBRE | Spain. Huelva, Corral de la Liebre. Doñana National Park. | 36.908 | -6.414 | 1.47 ± 0.02 (9) | 2x | SEV290229 |
| *Thymus mastichina subsp. donyanae* * | D08 | CORRALILLO OSCURO | Spain. Huelva. Corralillo Oscuro. Doñana National Park. | 36.987 | -6.414 | 1.51 ± 0.01 (8) | 2x | SEV290230 |
| *Thymus mastichina subsp. mastichina* * | M01 | CAZALLA | Spain. Sevilla, Cazalla de la Sierra. Sierra Morena de Sevilla Natural Park. | 37.908 | -5.802 | 2.64 ± 0.06 (8) | 4x | SEV290231 |
| *Thymus mastichina subsp. mastichina* | M02 | CORDOBA | Spain. Córdoba, El Vacar. ZEC Guadalmellato. | 38.100 | -4.833 | 1.58 ± 0.01 (9) | 2x | SEV290232 |
| *Thymus mastichina*<br>*subsp. mastichina* | M03 | SOUTH CARTAYA | Spain. Huelva, Cartaya. Campo Común de Abajo. | 37.238 | -7.080 | 1.56 ± 0.04 (8) | 2x | SEV290233 |
| *Thymus mastichina*<br>*subsp. mastichina* * | M04 | GILENA | Spain. Sevilla, Gilena. | 37.281 | -4.901 | 2.68 ± 0.02 (9) | 4x | SEV290234 |
| *Thymus mastichina*<br>*subsp. mastichina* * | M05 | GERENA | Spain. Sevilla, Gerena. | 37.561 | -6.159 | 2.71 ± 0.03 (11) | 4x | SEV290235 |
| *Thymus mastichina*<br>*subsp. mastichina* * | M06 | MONTE GORDO | Portugal. Algarve, Monte Gordo. | 37.186 | -7.488 | 2.78 ± 0.05 (9) | 4x | SEV290236 |
| *Thymus mastichina*<br>*subsp. mastichina* * | M07 | MADRID | Spain. Madrid. Hoyos de Manzanares. Cuenca Alta del Manzanares Regional Park. | 40.602 | -3.917 | 2.74 ± 0.04 (8) | 4x | SEV290237 |
| *Thymus mastichina*<br>*subsp. mastichina* * | M08 | VILLALUENGA | Spain. Cádiz, Villaluenga del Rosario. Sierra de Grazalema Natural Park. | 36.811 | -5.391 | 2.71 ± 0.07 (9) | 4x | SEV290238 |
| *Thymus mastichina*<br>*subsp. mastichina* * | M09 | RONDA | Spain. Málaga, Ronda, Pinsapar de la Nava. Sierra de las Nieves Natural Park. | 36.665 | -5.051 | 2.86 ± 0.06 (8) | 4x | SEV290239 |
| *Thymus mastichina*<br>*subsp. mastichina* * | M10 | NORTH CARTAYA | Spain. Huelva, Cartaya. Campo Común de Arriba. | 37.330 | -7.100 | 1.55 ± 0.02 (11) | 2x | SEV290240 |
| *Thymus mastichina*<br>*subsp. mastichina* * | M11 | EL PORTIL | Spain. Huelva, El Portil. | 37.203 | -7.014 | 1.57 ± 0.02 (12) | 2x | SEV290241 |
| *Thymus mastichina*<br>*subsp. mastichina* * | M12 | PAJARONCILLO | Spain. Cuenca, Pajaroncillo. | 39.930 | -1.726 | 2.73 ± 0.03 (6) | 4x | SEV290242 |
| *Thymus mastichina*<br>*subsp. mastichina* * | M13 | SAN CLEMENTE | Spain. Cuenca, San Clemente. | 39.338 | -2.472 | 2.84 ± 0.06 (8) | 4x | SEV290243 |
| *Thymus mastichina*<br>*subsp. mastichina* * | M14 | SERRA DA ESTRELA | Portugal. Guarda, Serra da Estrela Natural Park. | 40.379 | -7.507 | 1.49 ± 0.04 (7) | 2x | SEV290244 |
| *Thymus mastichina*<br>*subsp. mastichina* * | M15 | PONFERRADA | Spain. León, Ponferrada. | 42.564 | -6.771 | 2.64 ± 0.04 (6) | 4x | SEV290245 |
| *Thymus mastichina*<br>*subsp. mastichina* * | M16 | TORO | Spain. Zamora, Toro. | 41.516 | -5.414 | 2.7 ± 0.06 (7) | 4x | SEV290246 |
| *Thymus mastichina*<br>*subsp. mastichina* * | M17 | CERRO DA<br>CABEÇA | Portugal. Loulé, Cerro da Cabeça Alta. | 37.146 | -8.108 | 2.64 ± 0.04 (3) | 4x | SEV290492 |
| *Thymus mastichina*<br>*subsp. mastichina* * | M18 | PARIDERAS | Spain. Albacete, Parideras. | 38.582 | -2.526 | 2.52 ± 0.04 (8) | 4x | SEV290247 |
| *Thymus carnosus* | IB0432 | OUTGROUP 1 | Portugal. Faro. Ria Formosa Natural Park.<br>Praia da Anção. | 37.017 | -8.017 | N/A | N/A | MA887950 |
| *Thymus capitellatus* | IB0437 | OUTGROUP 2 | Portugal. Sesimbra. Aldeia do Meco. | 38.449 | -9.175 | N/A | N/A | MA643678 |
| *Thymus camphoratus* | IB0459 | OUTGROUP 3 | Portugal. Faro. Vilanova de Milfontes. | 37.740 | -8.788 | N/A | N/A | MA905261 |

### 2.2 Genome Size and Ploidy Level Assessments

Nuclear DNA amount was assessed from 344 individuals from the 40 studied populations. For these individuals, flow cytometry was performed either on fresh or silica-dried leaf material collected from the same plants used in the genomic workflow, or from seeds originating from those plants. *Lycopersicon esculentum* L. Stupické polní tyčkové rané cultivar (2C = 1.96 pg), and *Raphanus sativus* Saxa cultivar (2C = 1.11 pg) (Doležel *et al*. 1992), provided by the Centre of Plant Structural and Functional Genomics (Institute of Experimental Botany of the Czech Academy of Sciences), were used as internal standard. Leaf tissue of sample as well as internal standard were chopped together with a razor blade in a Petri dish in 1.2 ml of LB01 lysis buffer (Doležel *et al*. 1989), supplemented with Triton X-100 (8%; Girón *et al*. 2012) and 50 μg/ml RNase A, and stained with 50 μg/ml of propidium iodide. The nuclei suspension obtained was filtered through a 30 µm mesh CellTrics disposable filter (Partec GmbH, Munster, Germany), kept on ice for 5–30 min and measured in a flow cytometer CYTOMICS FC500-MP (Beckman Coulter, Fullerton, CA, USA) equipped with a 20 mW Ar-Ion laser at 488 nm. Acquisition was automatically stopped at 20,000 events. The total nuclear DNA content was calculated in each sample by multiplying the DNA content of the standard by the quotient between the 2C peak positions of the sample and the standard in the histogram of fluorescence intensities. As *Raphanus sativus* slightly but significantly overestimated genome size compared with *Lycopersicon esculentum* (t = -6.6302, df = 49; p < 0.01), we applied a correction factor of 0.95 to all measurements obtained with this standard, based on 50 samples analysed with both standards. Genome size assessments with both fresh material and dried leaves were highly correlated between the two methods (Deming Regression Fit, Pearson’s r = 0.99; N = 16) and that no significant differences were found between paired samples analysed with either method (Paired t-test, t = -0.19, df = 15, p = 0.853). Mean 2C values and standard deviations were calculated for each individual plant and for each population. Ploidy levels were assigned following Girón *et al*. (2012). Additionally, we estimated ploidy level from allelic ratios inferred from SNPs called from our genomic sequencing data (see section 2.4.4 below).

### 2.3 Molecular Protocols

#### 2.3.1 DNA Extraction and Genomic Library Preparation

DNA extraction was performed in eight individuals of each population. Between 15 and 20 mg of silica-dried foliar tissue per specimen was kept at −80°C for 15 min and then milled with a Mixer Mill MM301 (Retsch Inc., Haan, NRW, Germany). We used the DNeasy Plant Mini Kit (Qiagen, Hilden, NRW, Germany) and the GeneMATRIX DNA Purification Kit (EURx, Warsaw, Poland) for extraction of total cellular DNA, following manufacturer’s protocols. Precipitated DNA pellets were resuspended in Milli-Q ultrapure water (Merck KGaA, Darmstadt, HE, Germany). Relative DNA concentration (ng/µL) was measured with a dsDNA Quantification Assay Kit in a Qubit™ 3.0 fluorometer (ThermoFisher Scientific, Waltham, MA, USA). DNA extracts were sonicated with an ultrasonicator Bioruptor® Pico (Diagenode Inc., Liège, Belgium), for 10 cycles with 30’’ on and 30’’ off shearing times, to obtain an average fragment size of ∼250 bp. Dual-indexed genomic libraries, for Illumina®, were prepared using the NEBNext® Ultra™ II DNA Library Prep Kit and the NEBNext® Multiplex Oligos (Dual Index Primers Sets 1 and 2) from New England Biolabs (Ipswich, MA, USA), at half of the recommended volumes (Hale *et al*. 2020). Size selection was done with AMPure XP magnetic beads (Agencourt, Beverly, MA, USA) and the indexing PCR had 10 cycles in total. Concentration of all genomic libraries was checked using the Qubit® fluorometer, with 10% libraries being profiled for fragment distribution on a 4200 TapeStation System (Agilent Technologies, Santa Clara, CA, USA) with D5000 ScreenTape assays and reagents. Lastly, genomic libraries were normalised (10 nM) and pooled (∼24 libs/pool). Each pool contained ∼1500 ng DNA for an average fragment size of ∼290 bp.

#### 2.3.2 Hyb-Seq Implementation

Library pools were hybridised with the Angiosperms353 (Johnson *et al*. 2019) expert enrichment panel (Daicel Arbor Biosciences, Ann Arbor, MI, USA). A day-long hybridisation was carried out in an Applied Biosystems™ Veriti™ Thermal Cycler (Waltham, MA, USA), with settings 95°C 5 min, 65°C 5 min, and 65°C ∞. The hybridised, biotin-labelled baits were then captured with streptavidin-coated magnetic beads and further enriched with KAPA HiFi 2X HotStart ReadyMix PCR Kit (Roche, Basel, Switzerland), for 16 cycles, using the i5 and i7 forward and reverse “reamp” primers described in Meyer and Kircher (2010). Amplified PCR products were cleaned with Agencourt AMPure XP magnetic beads, quantified with a Qubit® fluorometer, and profiled on a 4200 TapeStation System. The enriched pools were then normalised and multiplexed, together with the plain genomic libraries, for a full Hyb-Seq implementation (Hale *et al*. 2020; Baker *et al*. 2022). Finally, this multiplexed Hyb-Seq pool (75% enriched library pools + 25% plain genomic libraries) was sequenced on an Illumina® NovaSeq X System producing 2 × 150 bp paired end reads at Macrogen Inc. (Seoul, South Korea).

### 2.4 Bioinformatic Analyses

#### 2.4.1 Data Mining, Target Assembly, and Data Matrix Refinement

Demultiplexed, raw sequencing reads were quality-checked with FastQC 0.12.1 (Andrews 2010) and MultiQC 1.25 (Ewels *et al*. 2016). Low-quality bases, short reads, and remaining adapters were filtered out using fastp 0.23.4 (Chen 2023). Nuclear targets and plastid coding DNA sequences (CDSs) were assembled with HybPiper 2.0.2 (Johnson *et al*. 2016), using the *mega353* target file (McLay *et al*. 2021). Here, we designed an ad hoc plastid CDS target file from the plastome of eight *Thymus* species available from an open access online repository (NCBI 1988). On average, out of ∼11M reads/individual, ∼1.18M reads/individual mapped to our 353 intended nuclear targets (i.e., ∼10.33% reads on target). Target success (measured as percentage of reads mapping to target loci in HybPiper) ranged between 4% and 54%, with ∼10.5% target success on average. Potential paralogy, checked with HybPiper, was not detected. Low-coverage (<⅓ median) samples and genes, as measured by the max_overlap.R script (see Shee *et al*. 2020), were removed. Finally, out of 323 individuals initially processed, we retained 304 accessions (301 from *Thymus* sect *Mastichina* plus 3 outgroups) and 332 supercontig sequences per accession, comprising both the exons and their flanking non-coding regions, for subsequent analyses. Detailed statistics from HybPiper and max_overlap.R can be found as a Zenodo repository (see Data Availability Statement below).

#### 2.4.2 Multiple Sequence Alignment (MSA) and Filtering

Balanced data matrices were aligned with MAFFT 7.520 (Katoh and Standley 2013), using the E-INS-i algorithm, and quality checked with AMAS (Borowiec 2016). Resulting MSAs were used to infer exploratory gene trees with FastTree 2.1.11 (Price *et al*. 2010). TreeShrink 1.2.1 (Mai and Mirarab 2018) was applied to automate removal of outlier branches from FastTree topologies and their corresponding MSAs. AMAS was used to select the threshold that maximised the proportion of parsimony-informative characters (P_PIC_), while minimising the percent of missing data. Target orthologs with P_PIC_ values <⅓ of the median were excluded. As a result, 301 individuals (298 from *Thymus* sect *Mastichina* plus 3 outgroups) and 329 supercontigs were retained for downstream analysis. Finally, the selected outlier-filtered MSAs were realigned again with MAFFT and further refined with trimAl 1.4.1 (Capella-Gutierrez *et al*. 2009). AMAS summaries can be found in the Zenodo repository (see Data Availability Statement).

#### 2.4.3 Gene and Species Tree Inference

Nuclear ML gene trees were estimated with IQ-TREE 2.2.0 (Minh *et al*. 2020). Selection of best-fit model of sequence evolution was done by ModelFinder Plus (Kalyaanamoorthy *et al*. 2017) and support values were calculated (1,000 bootstrap replicates) using both UFBoot2 (Hoang *et al*. 2018) and SH-aLRT (Anisimova *et al*. 2011) algorithms. Unsupported branch bipartitions (BS = 0) across ML gene trees were collapsed (see Mirarab 2019; Simmons and Gatesy 2021) using the Newick utilities 1.6.0 package (Junier and Zdobnov 2010). The species tree was estimated with ASTRAL III 5.7.8 (Zhang *et al*. 2018) under the multispecies coalescent (MSC) framework with branch support estimated as local posterior probabilities (LPP). Finally, the MSC species tree was scaled to present branch lengths in substitutions per site using RAxML-NG 1.2.2 (Kozlov *et al*. 2019). This scaled species tree was visualised using FigTree 1.4.4 (Rambaut 2018). As the plastome represents a canonical coalescent gene (c-gene; Doyle 2022), outlier-filtered, trimmed supercontig MSAs were concatenated using AMAS. Then, a plastid ML tree was inferred using IQ-TREE 2.2.0.

#### 2.4.4 SNP Calling

All *Thymus* sect*. Mastichina* accessions included in the phylogenomic analyses (298) were initially used for SNP calling, adapting the workflow designed by Slimp *et al*. (2021), with modifications (see Balant *et al*. 2025). Outgroups were excluded as this analysis was intended to examine genetic variation that segregates exclusively within the study populations. The variant detection was carried out with GATK4 4.5.0.0 (McKenna *et al*. 2010), using the longest (most-complete) supercontig sequence (assembled with HybPiper) for each of the 353 targets as a pseudo-genome reference. We opted for a conservative approach at SNP calling, by eliminating multi-allelic variants. A total of 82,137K single nucleotide polymorphisms (SNPs) were initially called. After a robust filtering, the SNP matrix was processed using BCFtools 1.20 (Danecek *et al*. 2021) and VCFtools 0.1.16 (Danecek *et al*. 2011). Linkage disequilibrium was avoided using PLINK 2.0 (Chang *et al*. 2015) with the command *--indep-pairwise* 50 5 0.2. Specifically, the command works within 50 variant sliding windows, by a step of 5 SNPs. Inside these windows, when two SNPs show r² correlation values higher than 0.2, one of them is removed. The final filtered SNP data matrix consisted of 254 individuals and 2,223 unlinked SNPs.

#### 2.4.5 Ploidy Level Estimation Based on Genomic Allelic Ratios

We estimated the ploidy level from our samples following the workflow designed by Viruel *et al*. (2019, 2023). Briefly, allelic frequencies were estimated with nQuire (Weiß *et al*. 2018). These frequencies were used to calculate allelic ratios and delta log-likelihood values per sample using the dplyr 1.1.4 (Wickham *et al*. 2023) and devEMF 4.5 packages (Johnson 2024) in R 4.3.2 (R Core Team 2022). Ploidy was then inferred using model-based approaches (Corrêa dos Santos *et al*. 2017), selecting the ploidy level corresponding to the model with the lowest log-likelihood value. These estimates were compared to our flow cytometry measurements (see section 2.2 above).

#### 2.4.6 Population Structure Analyses and Summary Statistics

Using the SNP matrix above, population structure analyses were carried out in ADMIXTURE 1.3.0 (Alexander *et al*. 2009; Alexander and Lange 2011), and the optimal number of clusters was estimated based on cross-validation (CV) error (metric that reflects the model’s predictive accuracy for different values of K). In parallel, STRUCTURE 2.3.4 (Pritchard *et al*. 2000), as implemented in the ipyrad toolkit 0.9.102 (Eaton and Overcast 2020), was also used to estimate the optimal number of genetic clusters. A range of K values (2–10) was tested in ten independent runs and the most likely number of clusters was selected by detecting the highest values of the ΔK statistic (Evanno *et al*. 2005). ADMIXTURE results for each K value were plotted using the pophelperShiny 2.1.1 R package (Francis 2017). R packages mapmixture 1.2.0 (Jenkins 2024) and ggplot2 (Wickham 2016) were used to obtain pie charts and project them onto a map obtained from QGIS 3.22 (QGIS.org 2024). Principal Component Analysis (PCA) and Discriminant Analysis of Principal Components (DAPC) were conducted in R package adegenet 1.3–1 (Jombart and Ahmed 2011). R packages poppr 2.9.6 (Kamvar *et al*. 2014) and ape 5.0 (Paradis and Schliep 2019) were also used to visualise the results.

The fixation index (F_ST_), inbreeding coefficient (F_IS_), and pairwise identity-by-descent (IBD) were calculated for each pair of the detected phylogenetic groupings and for each population using PLINK 2.0 (Hudson *et al*. 1992; Bhatia *et al*. 2013; Chang *et al*. 2015). Genetic variability within and among phylogenetic groupings was tested using analyses of molecular variance (AMOVA) with the poppr 2.9.6 R package. A Mantel test was performed using the vegan 2.6-8 R package (Oksanen *et al*. 2025), based on Pearson’s product-moment correlation. Nucleotide diversity (π) and Tajima’s D (Tajima 1989) were calculated using VCFtools. To identify private and shared SNPs among the phylogenetic groupings, we used the VennDiagram 1.7.3 R package (Chen and Boutros 2011). Analyses were performed using a random subsample of 11 individuals per grouping, corresponding to the sample size of the smallest grouping, in order to standardise grouping sizes and thus reduce bias in the number of private SNPs detected due to unequal sample size across groups.

#### 2.4.7 Introgression and Gene Flow Analyses

We reconstructed a Neighbour-Net phylogenetic network with SplitsTree6 (Huson and Bryant 2024) using both the concatenated gene matrix used for phylogenomic analyses and the filtered SNP matrix. This analysis allowed us to visualize the potential reticulation events among samples. Additionally, an ABBA-BABA test was performed among all genetic groupings using Dsuite 0.5 r58, following Malinsky *et al*. (2021), which relies on Patterson’s D statistic to estimate the genome-wide excess of shared derived alleles between two given groupings, in order to test for introgression events.

To further evaluate the presence of interploidy gene flow and to quantify its directionality, we applied fastsimcoal 2.8 (Excoffier *et al*. 2021), using Site Frequency Spectrum (SFS) files and the easySFS software (Gutenkunst *et al*. 2009). We compared four scenarios: (1) no gene flow (NOGF), (2) unidirectional gene flow from tetraploids to diploids (MIG21), or (3) from diploids to tetraploids (MIG12), and (4) bidirectional gene flow (MIGBI). We performed 50 runs, estimating parameters using ECM optimization cycles and coalescent simulations. The best-fitting model was selected based on the lowest median AIC value and verified using likelihood distributions. Finally, we used block-bootstrapping with 100 replicates to calculate confidence intervals for the parameter estimates, ensuring the reliability of our findings (Canty and Ripley 2024).

## 3 Results

### 3.1. Genome Size Measurements

DNA content estimated by flow cytometry revealed mean 2C values ranging from 1.32 to 3.07 pg (mean ± SD) in each analysed individual (Table S1). Mean 2C values in populations sampled by Girón *et al*. (2012) were on average 6% lower than those obtained in our study (Table 1), although those differences were low in terms of DNA amount (mean = 0.1 pg). Mean 2C values from sampled individuals exhibited a bimodal distribution: samples ranging from 1.32 to 1.84 pg were considered diploid, while those with mean 2C values from 2.31 to 3.07 pg were considered as tetraploid (Table S1), following Girón *et al*. (2012).

### 3.2 Phylogenomic Relationships in *Thymus* sect. *Mastichina*

The resulting species tree (Figures 1a and 2) presented *Thymus* sect. *Mastichina* divided into two phylogenetic groups (LPP = 0.78), respectively corresponding to diploid and tetraploid accessions. Following Morales (2010), the diploid group was composed of accessions identified as either *T. albicans* or *T. mastichina* subsp. *donyanae,* as well as accessions identified as *T. mastichina* subsp. *mastichina* in some populations (Table 1). The tetraploid group was composed of accessions identified as *T. mastichina* subsp. *mastichina*, distributed throughout the Iberian Peninsula, but with high variability in calyx length among and within individuals, also fitting the descriptions *T. mastichina* subsp. *donyanae* and *T. albicans* in relation to calyx length. The diploid group revealed two sister subgroups (LPP = 0.83). On one hand, the Hercynian subgroup, composed of diploid individuals phenotypically fitting *T. mastichina* subsp. *mastichina* from two geographically different areas—Serra da Estrela Natural Park (central Portugal; LPP = 0.98) and Córdoba (S Spain; LPP = 1)—and, on the other hand, the rest of the diploid populations, all of them located in SW Iberian Peninsula, that were further subdivided into three weakly-supported (LPP < 0.6), albeit geographically-structured subgroups (Figure 1a). The Algarve subgroup included individuals from the Algarve region (S Portugal), with phenotype fitting with *T. albicans*. The Doñana subgroup was mainly composed of individuals from Doñana National and Natural Parks (Huelva province, Andalusia, S Spain), with phenotype corresponding to *T. mastichina* subsp. *donyanae*, as well as nearby populations, some of which presented intermediate characters between *T. mastichina* subsp. *donyanae* and *T. albicans* while the westernmost populations of this group fitted with *T. mastichina* subsp *mastichina* (Table 1). Finally, the Cádiz subgroup mostly consisted of individuals from Cádiz province (Andalusia, S Spain) (Table S1). The plastome tree, based on 72 plastid CDSs showed a mixed distribution of individuals, with no clear groupings (Figure S1).

**FIGURE 1.**
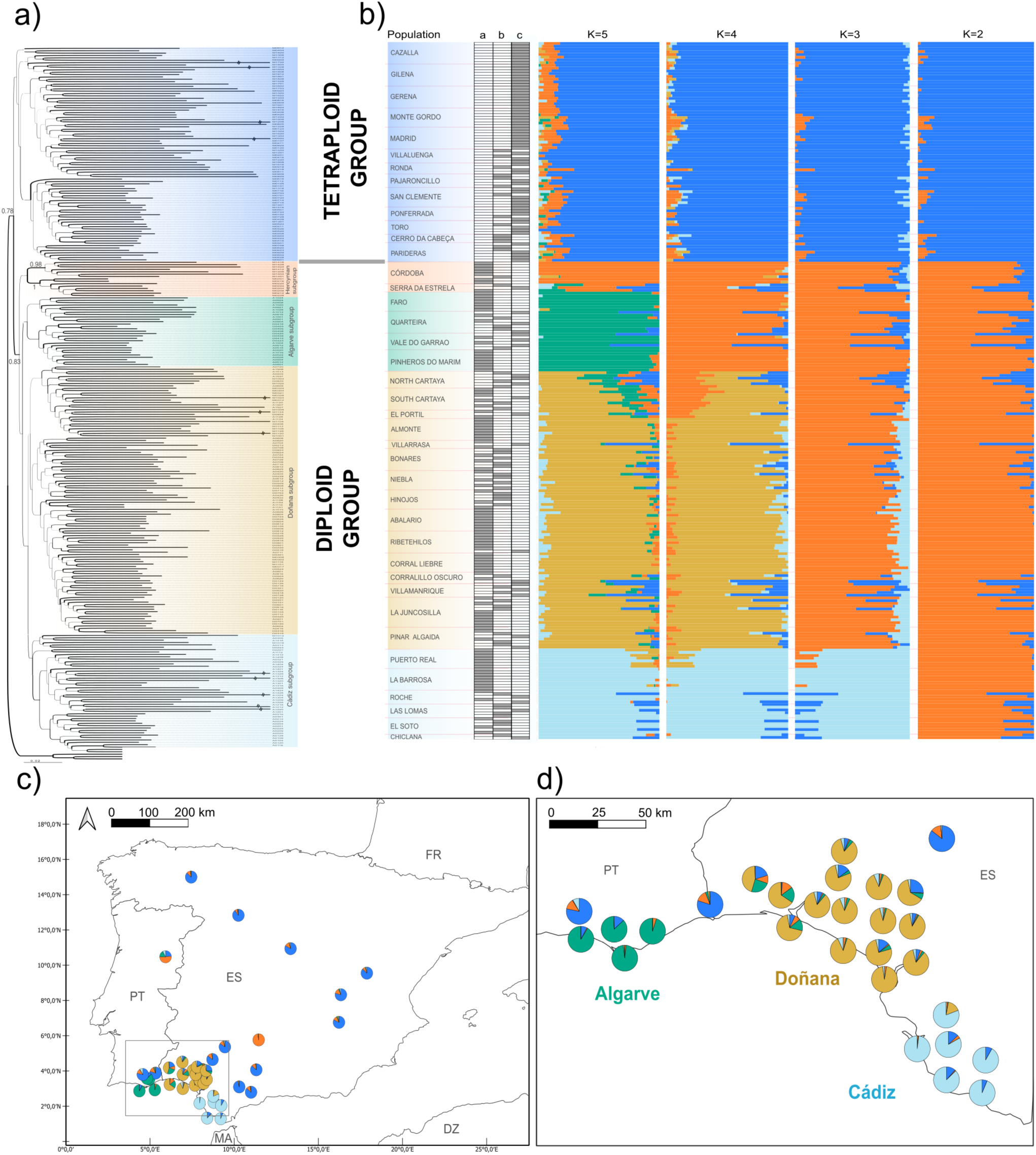
Phylogenomic relationships and population structure of Iberian endemic *Thymus* sect. *Mastichina* individuals. (a) Multispecies coalescent (MSC) tree obtained with ASTRAL-III (branch lengths in substitutions per site were scaled with RAxML-NG) for 329 nuclear orthologs. Support values, measured as local posterior probabilities (LPPs), under 75% not shown. (b) Population structure inferred with ADMIXTURE from 2,223 filtered and unlinked SNPs for the most likely clustering scenarios (K = 5 is the optimum; see Figure S2). Columns named “a”, “b” and “c” correspond to individual ploidy assignments (diploid, triploid and tetraploid, respectively) based on allelic ratio distributions as estimated with nQuire (see Figure S5). (c) Spatial distribution of genetic clusters across the Iberian Peninsula for *K* = 5. (d) An inset zooming into the SW Iberian Peninsula. Countries are denoted by their 2-digit ISO code.

**FIGURE 2.**
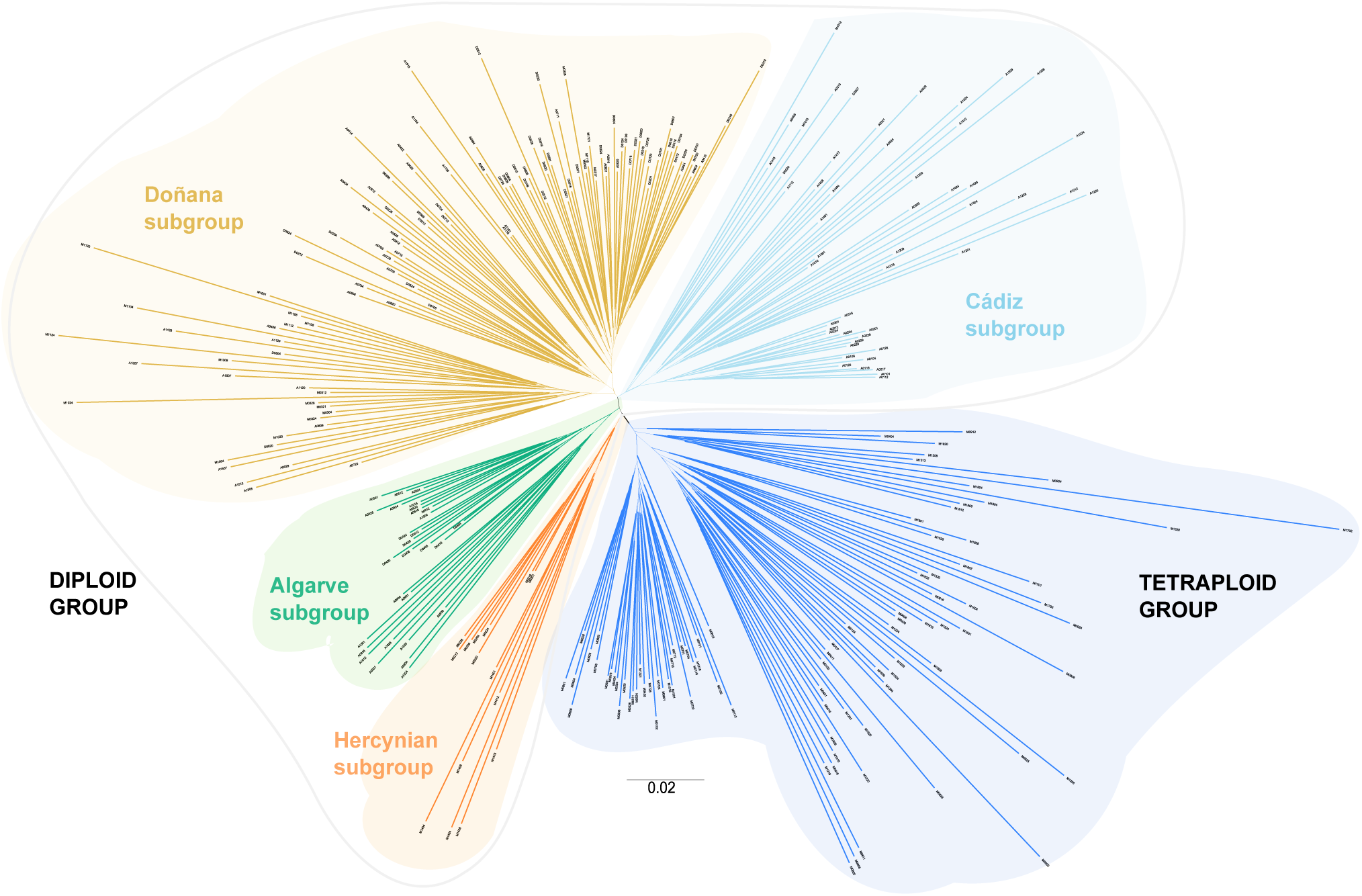
Unrooted MSC nuclear tree obtained with ASTRAL-III (branch lengths in substitutions per site were scaled with RAxML-NG) for 329 nuclear orthologs, showing the distribution of genetic groupings of *Thymus* sect. *Mastichina*. Colour scheme as in previous figure.

### 3.3 Ploidy Level Estimates from Allelic Ratios

Estimated allelic ratios per individual (Figure S2) ranged between 1 and 5.5, with median values from 1.24 to 2.67, in the diploid group, and from 1.77 to 3.04, in the tetraploid group. Model approaches classified the median allelic ratios per individual (Figure S2) into three levels: (a) low (1.24–1.80), (b) medium (1.70–2.25), and (c) high (1.80–3.04). These three allelic ratio levels fit the three ploidy models for which the nQuire pipeline specifically tests: diploid, triploid, and tetraploid, respectively. The tree categories were found in the diploid group and the two latter in the tetraploid group (Figure 1b). 2C DNA content (see section 3.1.) significantly differed among the three putative ploidy level estimates inferred from allelic ratio values (Kruskal-Wallis χ^2^ =84.47, df = 2, p < 0.001), although these differences were non-significant when considering only the diploid group (Kruskal-Wallis χ^2^ = 3.63, df = 2, F = 0.945, p = 0.163). Interestingly, the percentage of admixture of the tetraploid group into the diploid subgroups was significantly correlated with the mean allelic ratio per individual (Spearman rank correlation, L = 0.742; S = 226265, p < 0.001).

### 3.4 Phylogeographic Structure and Interploidy Admixture

The genetic structure of all sampled individuals is presented in Figure 1b. Optimal and suboptimal number of population structure groupings ranged between two and five, using the CV-error method, while the ΔK value supported that the optimal number of clusters was five (Figure S3). Population structure analysis for K = 2 distinguished tetraploid and diploid individuals into two distinct genetic groups. For K = 3 and K = 4, the tetraploid samples remained assigned to the same genetic group, and diploid samples were split into two and three genetic groupings, respectively. The likeliest five-cluster scheme (K = 5) was consistent with the phylogenetic groupings described above: the tetraploid *T. mastichina* group; and the diploid group, composed of the same four genetic subgroups (Algarve, Cádiz, Doñana, and Hercynian) observed in the phylogeny (Figure 1a,b), showing a distinct geographic pattern (Figure 1c,d). Interploidy admixture was observed across all K-schemes, with 78% of populations showing evidence of it in at least some individuals for K = 5 (Figure 1b).

The first five principal components of the PCA accounted for 94% of the variance and jointly supported the population structure for K = 5 (Figure 3a). Clustering analysis with DAPC (Figure 3b) was also coherent with the aforementioned five-cluster pattern, and largely consistent with the PCA. The two first linear discriminants (LD) of the DAPC explained >95% of cumulative variance. LD1 clearly separated tetraploid *vs.* diploid samples. LD1 and LD2 separated the four diploid subgroups.

**FIGURE 3.**
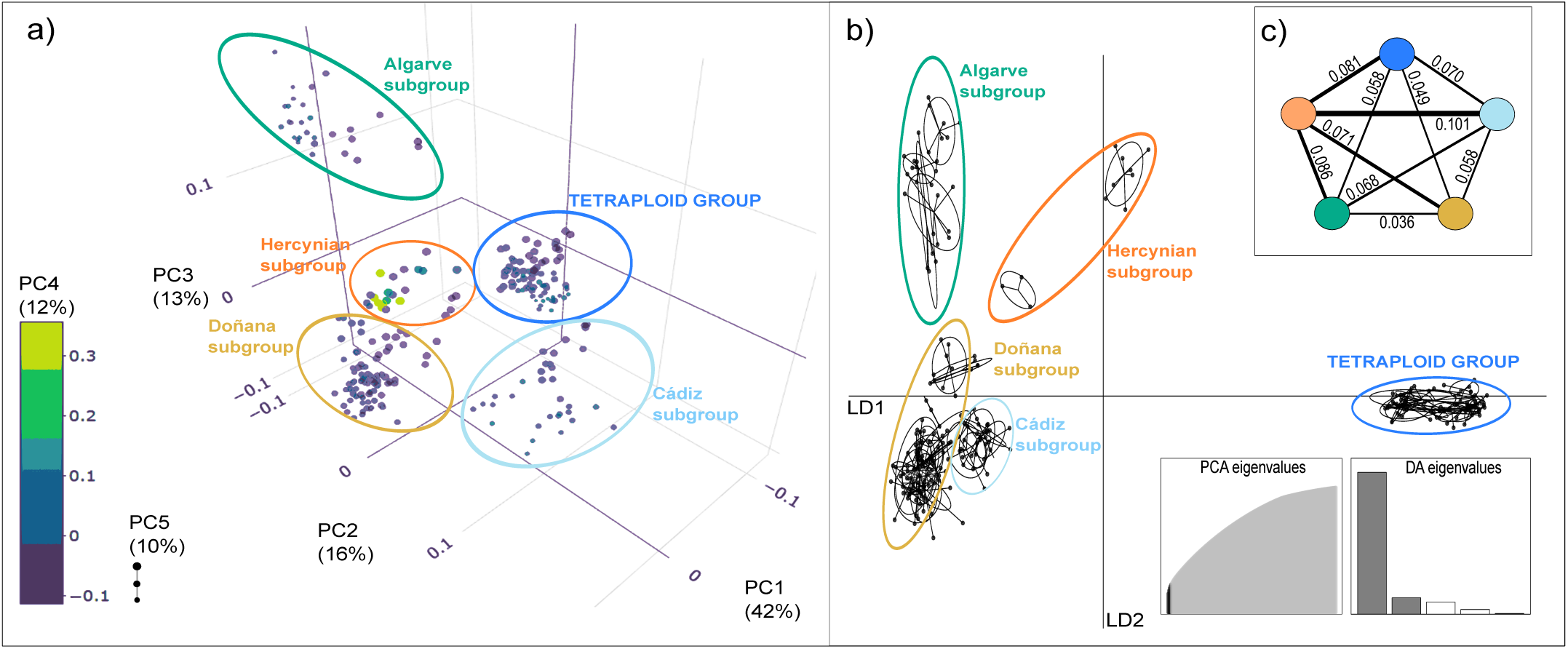
Graphical representation of a dimensionality reduction technique and a multivariate method for 254 samples based on 2,223 filtered and unlinked SNPs. (a) Five-dimensional Principal Component Analysis (PCA), and (b) Discriminant Analysis of Principal Components (DAPC). (c) Pentagonal diagram showing mean pairwise fixation indexes (line thickness proportional to F_ST_ value) between the five genetic groupings identified in the phylogenomic and population structure analyses (see Figures 1 and 2). Colour scheme as in previous figures.

Pairwise fixation indexes (F_ST_) indicated a moderate differentiation among the five identified genetic groupings (0.036 ≤ F_ST_ ≤ 0.101; Figure 3c), with the Hercynian subgroup showing the highest mean F_ST_, when compared with other subgroups (0.071 ≤ F_ST_ ≤ 0.101). In contrast, the Doñana subgroup showed the lowest FST (0.036 ≤ F_ST_ ≤ 0.071; Figure 3c). When comparing populations, the lowest F_ST_ index values were found within the tetraploid group and the diploid Doñana subgroup (Figure S4). The AMOVA test (Table S2) showed significant genetic structure in the four tested scenarios (K2-K5). The K2 clustering exhibited the highest levels of genetic variance among groups (28%; p < 0.001), followed by K3 and K5 (both 25%; p < 0.001).

The Mantel test showed a significant isolation-by-distance pattern among populations within the diploid genetic group (r = 0.83; p = 0.001), but no geographical isolation was detected among the more geographically spread populations of the tetraploid group (p = 0.161). This significant isolation-by-distance pattern was maintained when considering only the three diploid subgroups (Algarve, Doñana, and Cádiz) restricted to the SW Iberian Peninsula (r = 0.75; p = 0.001).

### 3.5 Genetic Diversity

Genetic diversity was analysed using different indicators. The mean inbreeding coefficient (F_IS_) per genetic grouping (Figure S5 and Table S3) ranged between −0.278 and 0.222. Tetraploid *T. mastichina* individuals showed the lowest mean inbreeding values (F_IS_ = −0.278), while diploid individuals showed in general higher inbreeding values (0.003 ≤ F_IS_ ≤ 0.222), especially the Hercynian subgroup (Table S3). The highest mean proportion of identical-by-descent (IBD) alleles was found in the tetraploid *T. mastichina* group (IBD = 0.223), while the lowest values were found in the diploid Cádiz subgroup (IBD = 0.055; Table S3). The nucleotide diversity (π) index showed, in general, low values (0.0012 ≤ π ≤ 0.0025; Figure 4a, Table S4); tetraploid populations had the highest mean π values (0.0015 < π < 0.0025; Figure 4a, Table S4). Tajima’s D values (Figure 4b, Table S4), ranged between −0.667 and 0.440, negative mean values were found in all populations within the Doñana subgroup (Figure 4b, Table S4). Private and shared SNPs in a balanced random subsample of analysed individuals showed a skewed pattern of SNP distribution among genetic groupings (Figure 5). On one hand, the tetraploid *T. mastichina* group shared more than 50% of the analysed SNPs with all the diploid subgroups (Figure 5). On the other, 44 private SNPs (2%) were detected in the tetraploid *T. mastichina* group, while in the diploid subgroups the number of private SNPs was much lower (3–11 SNPs/subgroup). The number of shared SNPs among groups and subgroups was also variable (Figure 5). The highest value of shared SNPs corresponded to the tetraploid group and the diploid group minus the Hercynian subgroup (172 SNPs; 8%), the second highest value was found for the tetraploid group and the diploid group excluding Cádiz subgroup (95 SNPs; 4%).

**FIGURE 4.**
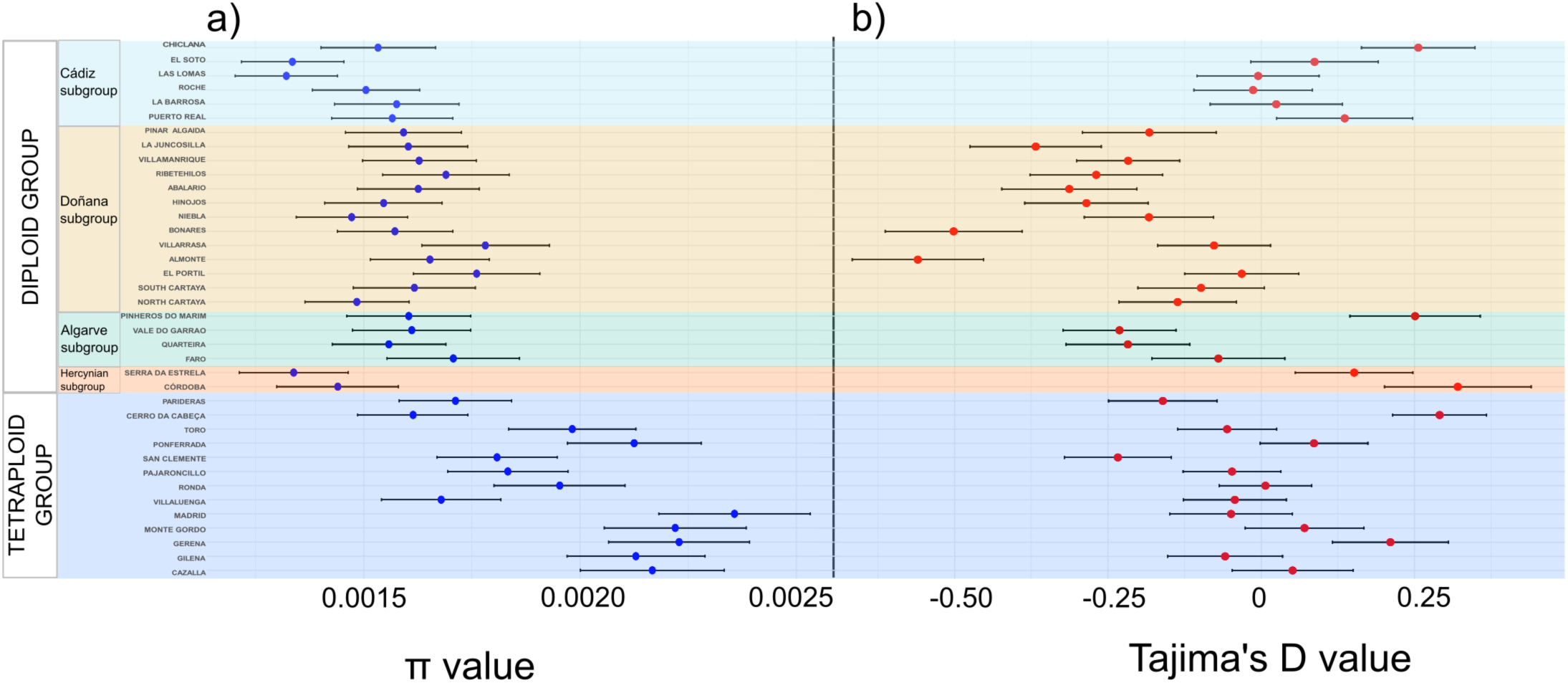
Genetic diversity estimated as (a) nucleotide diversity (π) and (b) Tajima’s D for *Thymus* sect. *Mastichina* populations (see Table S3). Colours denote the five genetic groupings identified in the phylogenomic and population structure analyses (see Figure 1). Colour scheme as in previous figures.

**FIGURE 5.**
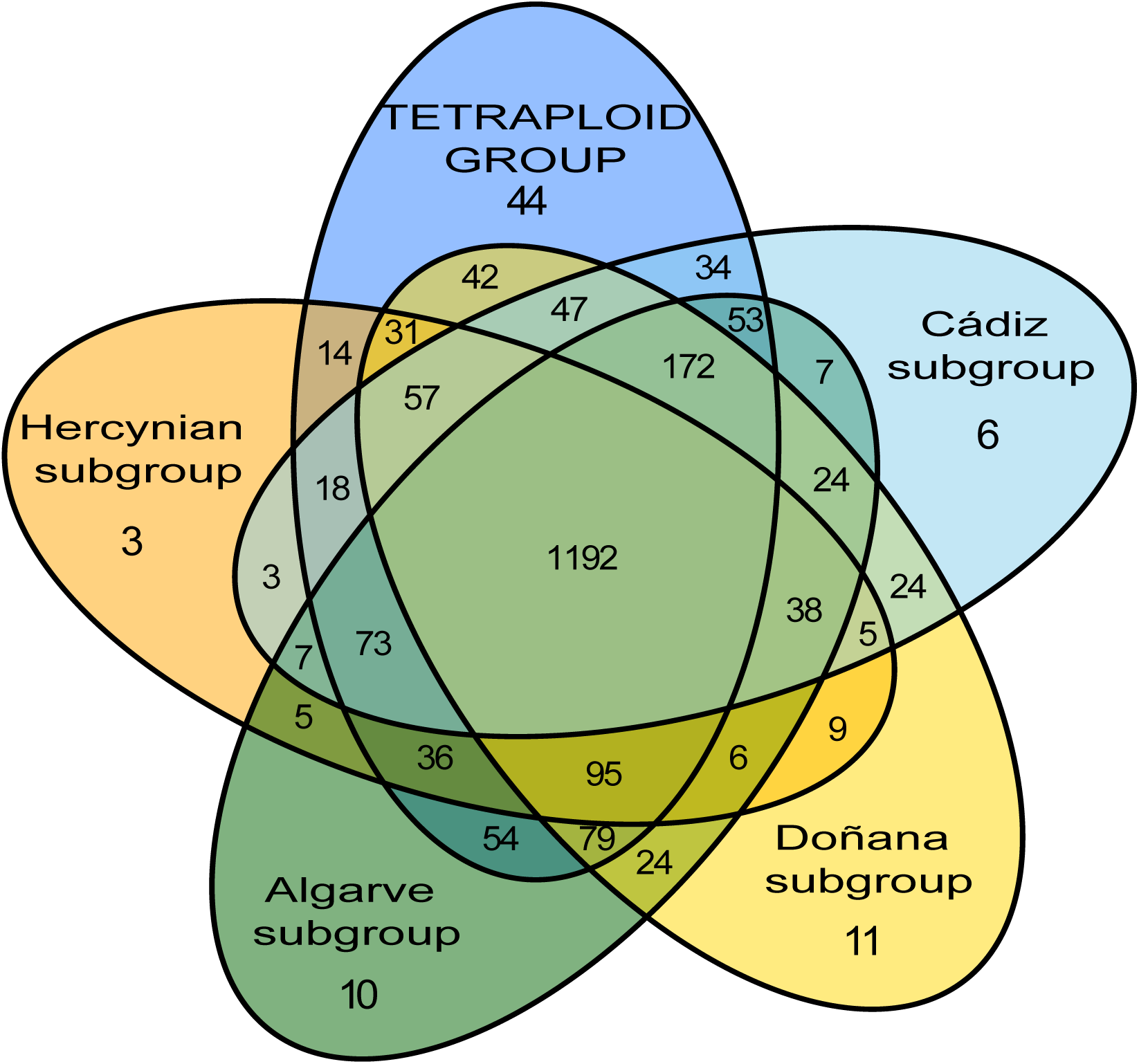
Venn diagram showing private SNPs (numbers in the ellipses without any overlap) and shared SNPs (numbers in the overlapping areas among ellipses) of a random subsample of 11 individuals for each genetic grouping within *Thymus* sect. *Mastichina*.

### 3.6 Reticulation, Introgression and Gene Flow Directionality

The Neighbour-Net phylogenetic network analysis provided a detailed view of the reticulations among the studied samples (Figure S6). Overall, the genetic structure of tetraploid and diploid groups was well defined; however, a dense reticulate core was observed in both networks for the concatenated supercontigs and SNP datasets. A high level of phylogenetic conflict was particularly evident among the diploid samples. Additionally, several reticulate connections were detected between the tetraploid clade and multiple diploid groups, especially the Doñana and Hercynian subgroups, as well as with the central core of the network.

The ABBA-BABA test provided evidence of introgression between the tetraploid *T. mastichina* group and the diploid Hercynian subgroup (ZLscore > 5.04; p < 0.0001; Figure 6 and Table S5), also supported by the high values of f4-ratio (Table S5) and F-branch statistic, which had a value of 1. A lower but significant introgression was also detected between the tetraploid *T. mastichina* group and the Doñana (Z-score = 2.76; p < 0.005) and Cádiz (Z-score = 2.349, p = 0.019) subgroups. Additionally, the diploid Algarve and Hercynian subgroups also showed significant introgression levels (Figure 6; Table S5).

**FIGURE 6.**
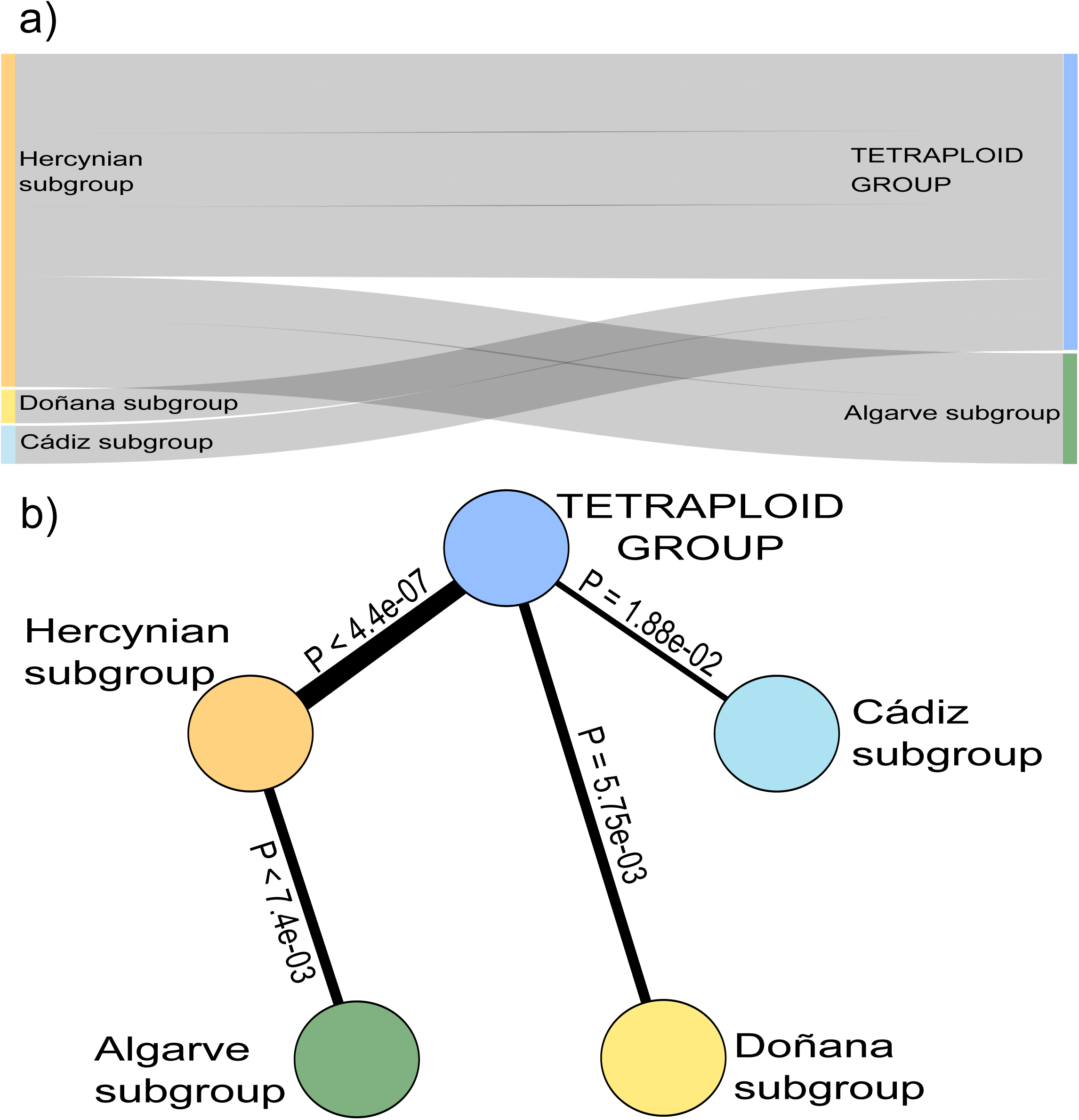
Diagrams showing the level of introgression as revealed by Dsuite (ABBA-BABA test, see Table S4) among the five genetic groupings in *Thymus* sect. *Mastichina*. (a) Sankey diagram illustrating gene flow between genetic groupings. (b) Schematic representation of introgression flow, with p-values indicating statistical significance. In both cases, the width of the bands or connection lines is proportional to the significance of the introgression. Colour scheme as in previous figures.

Coalescent simulations also detected interploidy gene flow in all tested scenarios (Figure 7, Table S6). Bidirectional gene flow was the best model for gene flow between tetraploids and the Algarve, Cádiz, and Doñana diploid subgroups, whereas unidirectional gene flow was the most plausible model from tetraploids to the Hercynian diploid subgroup. In all scenarios, overall migration intensity was consistently higher from tetraploids to diploids.

**FIGURE 7.**
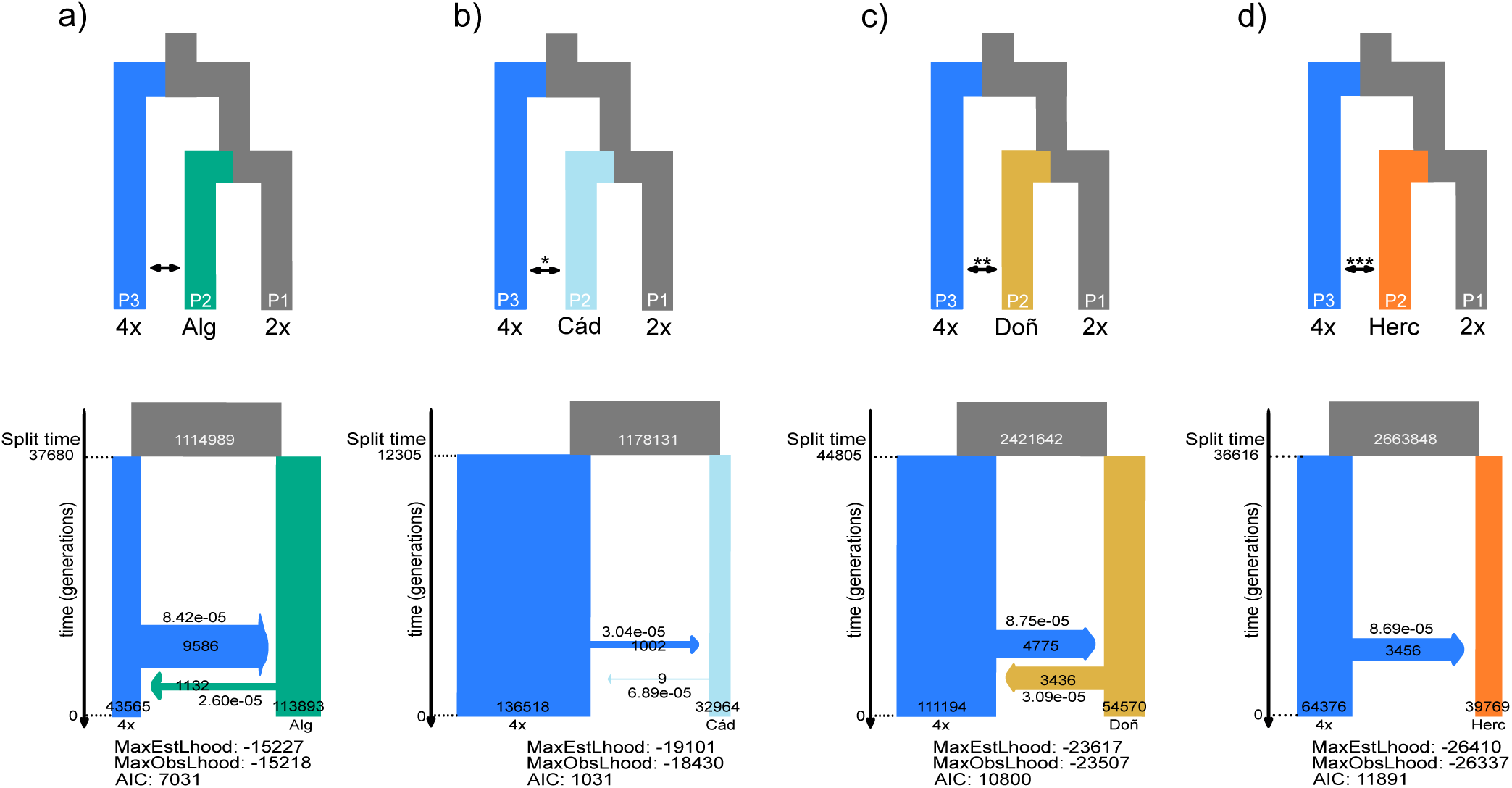
Strength and direction of interploidy introgression, as inferred with fastsimcoal2, among tetraploid and diploid groups of *Thymus* sect. *Mastichina*. Upper tree diagrams show ABBA-BABA tests of genome-wide interploidy introgression based on Patterson’s D statistic for the tetraploid group (4x) with each of the four diploid subgroups: (a) Algarve (Alg); (b) Cádiz (Cád); (c) Doñana (Doñ); and (d) Hercynian (Herc). 2x represents the remaining diploid subgroups (see Table S4). Significant tests are shown with asterisks (*p < 0.05; **p < 0.01; ***p < 0.001). Lower tree diagrams show preferred scenarios of isolation-with-migration as inferred from coalescent-based demographic analyses with fastsimcoal2. Bidirectional interploidy gene flow of asymmetrical strength between the tetraploid group and most diploid subgroups (a through c) and unidirectional gene flow from tetraploids to the Hercynian diploid subgroup (d). Migration is indicated forward in time. Columns and arrow thickness, and the number inside them, represent the effective population sizes and the number of migrants, respectively. Values above and below horizontal arrows represent migration rates. The maximum estimated log-likelihood (MaxEstLhood), the maximum observed log-likelihood (MaxObsLhood) and AIC index for each scenario are also shown.

## 4 Discussion

Our integrative analysis uncovers a complex evolutionary history in *Thymus* sect. *Mastichina*, with a clear diploid–tetraploid split and fine-scale phylogeographic structure among diploid populations, with four geographically defined diploid subgroups consistently recovered (Figures 1–3). Yet, plastid–nuclear discordance (Figures 1 and S1), allelic ratio variation (Figure S2), and evidence of gene flow—including interploidy introgression (Figures 6-7)—reveal dynamic and complex evolutionary processes. These patterns highlight ongoing diversification in this Iberian polyploid complex.

### 4.1 Ploidy Differentiation and Origin

Flow cytometry revealed a bimodal 2C DNA content in *Thymus* sect. *Mastichina*, matching diploid and tetraploid individuals (Girón *et al*., 2012). In parallel, the phylogenomic reconstruction confirmed that this section is composed of two sister clades—tetraploids vs. diploids—, the latter with four geographically-restricted subgroups (Algarve, Cádiz, Doñana, and Hercynian; Figures 1a and 2). SNP-based clustering methods (ADMIXTURE, PCA, DAPC) also supported this structure (Figures 1b and 3) and so did the Phylogenetic Neighbour-Net network (Figure S6). All tetraploids showed traits of *T. mastichina* subsp. *mastichina* (*sensu* Morales 2010), while diploids encompassed all section morphospecies: *T. mastichina* subsp. *mastichina* (in Hercynian and Doñana subgroups), *T. mastichina* subsp. *donyanae* (in Doñana), and *T. albicans* (in Algarve, Cádiz, and Doñana). Girón *et al*. (2012) also found diploids matching *T. mastichina* subsp. *mastichina* in Serra da Estrela (Hercynian) and Huelva (Doñana). Thus, the current taxonomy does not reflect the evolutionary history of the section.

Morales (1986) proposed that the tetraploid taxon originated from diploid progenitors in SW Iberian Peninsula and later colonised the entire peninsula, adapting to continental climates during Pleistocene cold phases. However, our results do not support this hypothesis. Additionally, he was unaware of *mastichina*-like diploid individuals in the Hercynian region (Morales, R., pers. com.). Our data reveal that most tetraploids share a percentage of genetic ancestry with the Hercynian diploids. Yet, Hercynian ancestry is not dominant, and *T. mastichina* appears as a genetically distinct lineage, not as a mixture. For that reason, we cannot establish a reliable hypothesis regarding the role of Hercynian diploids in the origin of the tetraploid group. The observed genetic pattern in the tetraploid group could reflect autopolyploidy from an extinct (ghost) diploid, followed by interploidy admixture (see Leal *et al*. 2024). Alternatively, an allopolyploid origin via hybridisation between extant Hercynian diploids and an unknown extant or ghost progenitor (Luo *et al*. 2017), followed by WGD and subgenome dominance could also be plausible (Bird *et al*. 2018; Liu *et al*. 2025). Further evidence, especially on potential diploid progenitors from other sections, is required to assess a putative allopolyploid origin of the tetraploid *T. mastichina*.

Polyploid species may exhibit higher allelic mutation rates across their multiple genomic complements, leading over time to greater genetic variation and, consequently, higher genetic diversity than diploids (Segarra-Moragues and Catalán 2003; Segarra-Moragues *et al*. 2004; Clo 2022; Campos *et al*. 2024). This enhanced variation may confer broader ecological tolerance, enabling polyploids to colonise new habitats more readily than diploids (Stebbins 1985; te Beest *et al*. 2012), and promoting diversification (Meudt *et al*. 2021). In *Thymus* sect. *Mastichina*, the widespread tetraploid *T. mastichina* group shows higher nucleotide diversity (π) than the geographically restricted diploid subgroups. Additionally, no isolation-by-distance pattern is detected in the tetraploid group, likely reflecting higher gene flow and connectivity among its populations. This is also suggested by the low F_ST_ and F_IS_ values in these populations, although we are aware that our study system does not meet all Hardy-Weinberg equilibrium assumptions (e.g., diploid populations and neutral genetic variation in all loci) and this could bias these estimates (Baudry and Depaulis 2003; McTavish and Hillis 2015; Liang and Dutheil 2025). Also, homeologous regions from ancient duplications may cause apparent excess heterozygosity and confound interpretations of variation and structure (Hohenlohe *et al*. 2010), our analyses are based on SNPs coming from nuclear orthologs (exons and their flanking regions), minimising such effects and ensuring more accurate estimates (Balant *et al*. 2025).

Alternatively, the reduced differentiation among distant tetraploid populations may reflect rapid range expansion after their emergence, likely due to a greater adaptive potential (Ehrendorfer 1980; HullLSanders *et al*. 2009; Treier *et al*. 2009; te Beest *et al*. 2012; Kiedrzyński *et al*. 2021). A plausible scenario involves diploid range contraction and habitat fragmentation (presumably *Quercus* forests; Valdés Castrillón *et al*. 1999), while tetraploids expanded their distribution by invading new habitats, as has been proposed for polyploids (Lumaret 1988; Kiedrzyński *et al*. 2021).

### 4.2 Diploid Lineage Structure and Phylogeographic Patterns

The phylogeographic structure observed in diploid populations of *Thymus* sect. *Mastichina* likely reflects a complex history of colonisation and diversification tied to the geological evolution of the Iberian Peninsula. The Hercynian subgroup, comprising two populations in the ancient Iberian Massif (Eguíluz *et al*. 2000), consistently appears as sister to the other diploid subgroups in both the phylogenomic tree and ADMIXTURE analyses (Figure 1). This group may represent a relict lineage that originated and persisted in geologically stable environments. Their higher F_IS_ and F_ST_ values point to low connectivity between populations and a marked population differentiation (Young *et al*. 1996; Charlesworth 2003). The absence of *Thymus* sect. *Mastichina* representatives in North Africa, despite floristic similarity across the Strait of Gibraltar (Rodríguez-Sánchez *et al*. 2008), supports its origin postdating the strait’s formation (∼5.3 Mya; Krijgsman *et al*. 1999). In fact, divergence time estimates for *Thymus* suggest a relatively recent origin at the Plio-Pleistocene boundary (∼2.58 Mya; Drew and Sytsma 2012). If *Thymus* section *Mastichina* originated somewhere in the Iberian Massif, a colonisation of the Algarve (a Mesozoic basin later restructured by the Alpine Orogeny; Ramos *et al*. 2020) by the most recent common ancestor (MRCA) of all diploid lineages, is plausible. Another likely migration could have occurred southward into the Cádiz lowlands, southwest of the Betic Mountains, formed in Alpine Orogeny (Vergés *et al*. 2019). In both cases, this southward dispersal into more meridional latitudes is a well-documented biogeographical pattern in plant lineages, linked to glacial–interglacial dynamics. In fact, Algarve and Cádiz are recognised as putative Western Mediterranean refugia (Gómez and Lunt 2007; Médail and Diadema 2009).

In contrast, the Doñana region, between Algarve and Cádiz, is geomorphologically younger. Its current landscape was shaped by Guadalquivir River sediments and sea-level rise (Rodríguez-Ramírez *et al*. 2019) with stabilization during the Holocene (Rodríguez Vidal 2005; Morales-Molino *et al*. 2011), making it one of the last regions within the current distribution of *Thymus* sect. *Mastichina* to become ecologically available. Negative Tajima’s D values in several Doñana populations may indicate a post-bottleneck expansion. The phylogenetic proximity with the Cádiz subgroup could point to a recent colonisation of Doñana’s sand dunes by the MRCA of these sister subgroups. However, the low branch support in our phylogeny limits inference on the colonisation source after Doñana dune formation. Overall, genomic and geological evidence suggests a scenario in which the diploid group originated inland (Iberian Massif) and expanded southwards to Algarve and Cádiz refugia and, more recently, to Doñana. Future work estimating divergence times and diversification rates will help test and refine this hypothesis.

Populations of the three diploid subgroups in SW Iberian Peninsula (Algarve, Cádiz, and Doñana) show clear genetic differentiation and significant west-east isolation-by-distance pattern, despite their adjacent distribution. Besides their colonization history, geographic features like the Guadiana and Guadalquivir riverbeds, between Algarve-Doñana and Doñana-Cádiz, respectively, could be acting as barriers to gene flow (see Ortiz *et al*. 2008). The absence of significant introgression among these adjacent groups (Table S5) aligns with their low gene flow. Also, the high inbreeding levels (Table S3), especially in the Cádiz subgroup, could indicate low functional connectivity among populations (Young *et al*. 1996; Charlesworth 2003). Interestingly, a population of the Doñana subgroup (Pinar Algaida) is currently located on the eastern margin of the Guadalquivir estuary (Cádiz province). However, its location likely reflects local coastal dynamics, not crossLriverbed dispersal. Around 6.5 Kya, a marine transgression (peaking at current sea level) filled the estuary with sediments, reshaping the shoreline into sandy barriers and marshes. Then, ∼5 Kya, an accelerated shoreline progradation built a promontory connecting the western margin of the ancient estuary. Later erosional pulses (e.g., storms, tsunamis) isolated this promontory as a small island and sculpted it into present-day Algaida, nowadays on the eastern estuary bank (Rodríguez-Ramírez 2008).

### 4.3 Reticulate Evolution, Admixture and Gene Flow

We found patterns among diploid and tetraploid lineages that point to a complex evolutionary history on *Thymus* sect. *Mastichina*. The observed shared heterozygosity in ADMIXTURE analysis (Figure 1b) and the evidence of reticulate evolution observed in Neighbour-Net networks (Figure S6). Such a structure is consistent with ILS following recent or rapid divergence events, especially among diploid lineages. Additionally, signals of interploidy gene flow between diploid and tetraploid lineages of *Thymus* sect. *Mastichina* provide evidence for secondary contact between ploidy levels throughout the evolutionary history of the group. ABBA-BABA tests identified significant introgression from tetraploid *T. mastichina* into several diploid subgroups—especially the Hercynian, but also Doñana and Cádiz (Figure 6). Coalescent-based analyses (Figure 7) also detected consistent patterns of interploidy gene flow across ploidy levels in all diploid subgroups. Although bidirectional gene flow emerged as the best-supported model between tetraploids and most diploid groups, the overall intensity of gene flow was consistently higher from tetraploids to diploids, pointing to asymmetric interploidy dynamics. This pattern is further supported by allele frequency distributions (Figure S2), allele-ratio-based ploidy inference, SNP-based ancestry proportions (Figures 1 and 3), and the Neighbour-Net network (Figure S6). Altogether, these results suggest recurrent and geographically widespread gene flow between ploidy levels, particularly in contact zones.

The clear bimodal distribution of 2C DNA content in *Thymus* sect. *Mastichina* supports a diploid–tetraploid distinction, yet our allele-ratio-based ploidy model also identifies triploid and tetraploid profiles within the diploid cytometric group. This mismatch suggests that this approach (Viruel *et al*. 2019, 2023), relying on nQuire (Weiß *et al*. 2018) and ploidyNGS (Corrêa dos Santos *et al*. 2017), may not be fully reliable in polyploid complexes with known or suspected interploidy admixture, as no 2C DNA content differences were found among inferred ploidy categories. Other allele-ratio-based models, such as gbs2ploidy (Gompert and Mock 2017), nQuack (Gaynor *et al*. 2024), or vcf2ploidy (DeWaters 2020), could face similar limitations. However, it is important to note that these models are not fully equivalent to nQuire, as they rely on different mathematical assumptions, and their performance would need to be tested empirically to assess how they behave in cases of interploidy admixture. Rather than reflecting current ploidy levels, the triploid and tetraploid categories inferred by the Viruel *et al*. (2019, 2023) workflow may reflect past or ongoing introgression (Nieto Feliner *et al*. 2020; Wang *et al*. 2023; Brown *et al*. 2024), with the fixation of homeolog loci from tetraploid genomes into diploid lineages. This hypothesis is further supported by significant correlations between the frequency assignment of diploidy to the tetraploid genetic group and allele ratios, and by introgression signals detected through the ABBA-BABA test, which revealed gene flow between tetraploid *T. mastichina* and the diploid Cádiz, Doñana, and Hercynian subgroups. The observed pattern could suggest not only recent but also ancient episodes of introgression, potentially dating back to periods soon after the tetraploid emergence.

Historical gene flow is likely in contact zones where diploids and tetraploids have long coexisted, enabling hybridisation and backcrossing. Over time, introgressed alleles may become fixed in recipient genomes, leaving persistent genomic signatures that may not necessarily reflect active gene exchange (Soltis and Soltis 2009; Nieto Feliner *et al*. 2020; Stull *et al*. 2023). This could explain interploidy admixture in isolated diploid populations such as Serra da Estrela (Hercynian subgroup). In contact zones of diploid and tetraploid populations, ongoing low-level gene exchange, possibly mediated by triploid bridges, could be plausible. Our coalescent-based demographic analyses, indicating gene flow mainly from tetraploids into diploids, also support this hypothesis. While the extent of recent interploid admixture remains uncertain, our results suggest that historical hybridisation and subsequent backcrossing have left a lasting imprint on the genomic architecture of several diploid populations (e.g., North Cartaya, Villamanrique, Serra da Estrela). Diploid individuals with traits from different morphospecies further suggest repeated interploid admixture and complex introgressed genotypes. These patterns align with reticulate evolution, where diploid–tetraploid barriers are porous under certain spatial-ecological conditions (Kauai *et al*. 2024; Bartolić *et al*. 2024, 2025). Introgressed alleles from tetraploids may be fixed or remain at intermediate frequencies in diploids, increasing heterogeneity and blurring taxon boundaries. The overall genomic cohesion of tetraploid *T. mastichina* may suggest limited recent diploid introgression, also detected by the coalescent-based demographic modelling, supporting asymmetrical, context-dependent interploid admixture mediated by population density, ecological compatibility, and hybrid fertility (VallejoLMarín *et al*. 2016; Kauai *et al*. 2024; Leal *et al*. 2024; Bartolić *et al*. 2024, 2025).

### 4.4 Evolutionary and Conservation Implications

Our results offer valuable insights and serve as a guiding example for the conservation of taxonomically complex or cryptic species (Bickford *et al*. 2007; Fišer *et al*. 2018; Hending 2025), where morphology alone proves inadequate (e.g., phenotypic plasticity; Matesanz *et al*. 2010). In *Thymus albicans* s.l. (referring to all diploids in *Thymus* sect. *Mastichina*), morphospecies poorly reflect underlying biological lineages. We recommend flow cytometry in conservation assessments to identify and monitor cytotypes and genetic structure (Heslop-Harrison *et al*. 2023). This rapid, cost-effective method can be easily integrated into routine prospections of new populations for conservation purposes. Given growing threats from tourism, coastal development, and land-use change (Valdés Castrillón *et al*. 1999; Cabezudo *et al*. 2005; Morales 2010), conservation should prioritise the four genetic subgroups identified within *T. albicans* as distinct units, as each lineage preserves the species’ genetic and ecological legacy.

Low genetic diversity, high inbreeding, and isolation—especially in Hercynian and Cádiz populations—suggest reduced gene flow and potential reproductive barriers, which may compromise long-term population viability (Young *et al*. 1996; Charlesworth 2003; Campos *et al*. 2024). This is especially relevant in gynodioecious species like the endangered *T. albicans*, where the effects of such cross-fertilisation barriers may be exacerbated in populations with female-biased sex ratios, as observed in many *Thymus* species (Karbstein *et al*. 2019) and also many of the studied populations (Berjano, pers. obs.). Future conservation-oriented analyses should incorporate additional variables, such as ecological features, population threat levels, and protection status, to provide more specific management recommendations.

## 5 Conclusions

Our integrative genomic approach confirms that *Thymus* sect. *Mastichina* comprises two sister groups with distinct ploidy levels. The tetraploid *T. mastichina* likely arose from related diploids, and shows no genetic structuring, consistent with recent expansion. In contrast, diploid populations exhibit complex structuration shaped by historical geomorphology, isolation, ILS and occasional interploidy introgression. Despite genetic divergence, phenotypic overlap and signs of reticulate evolution challenge current taxonomy, revealing cryptic diversity and underscoring the need for an integrative taxonomic framework combining morphological, cytogenetic, genomic, and geographic data. We advocate a taxonomic re-evaluation using multi-source evidence and emphasise the urgent conservation of the distinct diploid lineages of the endangered *T. albicans* s.l.—including not only the *‘albicans’* morphotype (Algarve, Cádiz) but also the ‘*mastichina*’ and ‘*donyanae*’ diploid morphotypes of the Hercynian and Doñana subgroups—, which represent unique evolutionary trajectories and may harbour critical adaptive potential. Future work should expand flow cytometry sampling across the Hercynian range and reassess morphological distinctions between diploid and tetraploid ‘*mastichina*’ morphotypes and select the most relevant populations for the effective conservation of the lineages. Additionally, estimates of divergence times, diversification rates, and phylogenetic networks are needed to fully resolve the evolutionary history of these ecologically and economically valuable Iberian taxa.

## Supporting information

Suppplemental Table S1

Suppplemental Table S2

Suppplemental Table S3

Suppplemental Table S4

Suppplemental Table S5

Suppplemental Table S6

Suppplemental Figure S1

Suppplemental Figure S2

Suppplemental Figure S3

Suppplemental Figure S4

Suppplemental Figure S5

Suppplemental Figure S6

## Author Contributions

R.B., D.N., and M.d.l.E. conceived the project, designed the methodological approach, and secured funding. DD, R.B., J.C.d.V., M.A.O., D.N.L., and F.J.G.-C. carried out the field work. L.P. designed the molecular and computational workflows, assisted by R.B. M.A.O., and F.J.G.-C. R. B., D.D, J.C.d.V., F.J.G-C., and L.P., conducted molecular work. R. B., D.D, J.C.d.V., F.J.G-C and M.A.O conducted flow cytometry analyses. F.J.G.-C., R.B., M.A.O. and L.P. analysed the data. R.B. and L.P. supervised the study. F.J.G.-C., R.B, L.P., and M.A.O. drafted the manuscript. All authors reviewed and edited subsequent versions and approved of the final one.

## Acknowledgements

This study is part of the Conserva3 project (TED2021-130133B-I00) funded by MCIN/AEI/10.13039/ 501100011033 with funds from the European Union “NextGenerationEU”/PRTR. LP benefited from a Ramón y Cajal grant (RYC2021-034942-I) funded by MCIN/AEI/10.13039/501100011033 and the European Union “NextGenerationEU”/PRTR.

Collection permits were obtained from Consejería de Sostenibilidad, Medio Ambiente y Economía Azul, Junta de Andalucía (Spain), Doñana National Park, and from Departamento de Conservação da Natureza e Biodiversidade (Portugal, permits no. 576–580/REC/2023). In Doñana National Park, logistic and technical support was provided by ICTS-RBD-CSIC, Ministry of Science and Innovation, and co-financed by FEDER Funds.

Authors would like to thank Javier Jiménez-Lopez, Javier López Tirado, Billy James Williams, Esther Martín Carretié, Andrés Melero, as well as the staff of the Environmental Agency and the Botanical Network of the Andalusian Government (Junta de Andalucía) and of the Doñana National Park for their support in population location and sample collection. Ricarda Riina provided DNA templates for *Thymus* outgroups.

Genomic libraries were prepared at the DNA lab of the Herbarium Research Services (CITIUS, University of Seville). Flow cytometry measurements were performed at the facilities of the Biology Research Services (CITIUS, University of Seville). Part of the costs of those Research Services were funded by 2023 and 2024 calls of VII PPIT of the University of Sevilla. Computational workflows were implemented in the Hercules supercomputer at the Scientific Computer Centre of Andalucia (CICA). Additional thanks go to Manica Balant, Matt Johnson, Rafa Albadalejo, Xavier Picó, Isaac Overcast, Paolo Bartolić, Marcial Escudero, and Ana Valdés, for their analytical advice, and to four anonymous reviewers that helped improve the manuscript.

## Data Availability Statement

The raw Hyb-Seq data are available on NCBI Sequence Read Archive (SRA) under BioProject PRJNA1326772. The code pertaining to the implemented analyses, the ad hoc plastid CDS target file, output datasets, and spreadsheets with summary statistics are available through Zenodo repository 10.5281/zenodo.15331851.

