## Supplementary material for "Patterns of Interploidy Admixture in Polyploid Complexes: Insights from *Thymus* sect. *Mastichina* (Lamiaceae)": Suppplemental Table S2

**Table S2**. AMOVA results

| **K** | **Source** | **df** | **SSD** | **MS** | **Est.Var** | **%** | **Phi** |
| --- | --- | --- | --- | --- | --- | --- | --- |
| K2 | Between groups | 1 | 4435 | 4435 | 38.80 | **28** | 0.2829 |
|  | Within groups | 252 | 24775 | 98 | 98.34 | **72** |  |
|  | Total | 253 | 29210 | 115 | 137.11 | 100 |  |
| K3 | Between groups | 2 | 4922 | 2461 | 31.85 | **25** | 0.2427 |
|  | Within groups | 251 | 24288 | 97 | 96.76 | **75** |  |
|  | Total | 253 | 29210 | 115 | 128.62 | 100 |  |
| K4 | Between groups | 3 | 5540 | 1847 | 29.40 | **24** | 0.2368 |
|  | Within groups | 250 | 23670 | 95 | 94.68 | **76** |  |
|  | Total | 253 | 29210 | 115 | 124.73 | 100 |  |
| K5 | Between groups | 4 | 5907 | 1477 | 30.59 | **25** | 0.2463 |
|  | Within groups | 249 | 23303 | 94 | 93.58 | **75** |  |
|  | Total | 253 | 29209 | 115 | 124.17 | 100 |  |

df = degrees of freedom; SSD = sum of squared deviation; MS = mean square; Est. Var. = estimated variance; % = percentage of total variance; Phi = genetic differentiation coefficient. Significance test P<0.001 are in bold
