## Supplementary material for "Patterns of Interploidy Admixture in Polyploid Complexes: Insights from *Thymus* sect. *Mastichina* (Lamiaceae)": Suppplemental Table S3

**Table S2**. Pairwise fixation index (Hudson F_ST_), inbreeding coefficient (F_IS_), and identity-by-descent (IBD) values in each studied population.

| **Genetic group** | **Population** | **F_ST_** | **F_IS_** | **IBD** |
| --- | --- | --- | --- | --- |
| TETRAPLOID  GROUP | MONTE GORDO | 0.007 | -0.495 | 0.334 |
|  | PARIDERAS | 0.052 | -0.039 | 0.099 |
|  | VILLALUENGA | 0.063 | 0.0021 | 0.061 |
|  | PAJARONCILLO | 0.041 | -0.1494 | 0.152 |
|  | SAN CLEMENTE | 0.037 | -0.1457 | 0.186 |
|  | PONFERRADA | 0.034 | -0.3503 | 0.247 |
|  | TORO | 0.052 | -0.2360 | 0.172 |
|  | MADRID | 0.029 | -0.5690 | 0.347 |
|  | RONDA | 0.017 | -0.2435 | 0.181 |
|  | CERRO DA CABEÇA | 0.062 | 0.0138 | 0.106 |
|  | CAZALLA | 0.022 | -0.4481 | 0.339 |
|  | GERENA | 0.038 | -0.5155 | 0.362 |
|  | GILENA | 0.035 | -0.4017 | 0.323 |
| **Mean** |  | **0.036** | **-0.2785** | **0.223** |
| Hercynian  subgroup | CÓRDOBA | 0.112 | 0.1529 | 0.101 |
|  | SERRA DA ESTRELA | 0.116 | 0.2914 | 0.029 |
| **Mean** |  | **0.114** | **0.2222** | **0.065** |
| Algarve  subgroup | FARO | 0.069 | -0.0633 | 0.120 |
|  | PINHEIROS DO MARIM | 0.088 | 0.0080 | 0.126 |
|  | QUARTEIRA | 0.071 | 0.0629 | 0.056 |
|  | VALE DO GARRAO | 0.064 | 0.0066 | 0.075 |
| **Mean** |  | **0.072** | **0.0035** | **0.094** |
| Doñana  subgroup | PINAR ALGAIDA | 0.055 | 0.0470 | 0.103 |
|  | ABALARIO | 0.036 | -0.0120 | 0.173 |
|  | CORRALILLO OSCURO | 0.039 | 0.0096 | 0.112 |
|  | ALMONTE | 0.040 | -0.0369 | 0.186 |
|  | CORRAL LIEBRE | 0.051 | -0.0388 | 0.163 |
|  | BONARES | 0.046 | 0.0805 | 0.087 |
|  | NORTH CARTAYA | 0.047 | 0.0922 | 0.034 |
|  | SOUTH CARTAYA | 0.054 | -0.0016 | 0.155 |
|  | HINOJOS | 0.049 | 0.0887 | 0.126 |
|  | NIEBLA | 0.064 | 0.1157 | 0.069 |
|  | EL PORTIL | 0.043 | 0.0061 | 0.082 |
|  | RIBETEHILOS | 0.055 | -0.0555 | 0.188 |
|  | VILLARRASA | 0.044 | -0.0551 | 0.151 |
|  | VILLAMANRIQUE | 0.028 | 0.0325 | 0.074 |
|  | LA JUNCOSILLA | 0.066 | -0.0261 | 0.133 |
| **Mean** |  | **0.047** | **0.0164** | **0.122** |
| Cádiz  subgroup | LA BARROSA | 0.074 | 0.0210 | 0.101 |
|  | CHICLANA | 0.091 | 0.2335 | 0.030 |
|  | LAS LOMAS | 0.100 | 0.1946 | 0.025 |
|  | PUERTO REAL | 0.086 | 0.0306 | 0.091 |
|  | ROCHE | 0.068 | 0.0702 | 0.057 |
|  | EL SOTO | 0.102 | 0.1916 | 0.029 |
| **Mean** |  | **0.086** | **0.1236** | **0.055** |
