## Supplementary material for "Patterns of Interploidy Admixture in Polyploid Complexes: Insights from *Thymus* sect. *Mastichina* (Lamiaceae)": Suppplemental Table S4

**Table S4**. Nucleotide diversity (**π**) and Tajima’s D values for each genetic grouping and population.

| **Genetic grouping** | **Population** | **Nucleotide diversity (π)** | | | **Tajima D** | | |
| --- | --- | --- | --- | --- | --- | --- | --- |
|  |  | **mean** | **min** | **max** | **mean** | **min** | **max** |
| TETRAPLOID  GROUP | MONTE GORDO | 0.0022 | 0.0021 | 0.0024 | 0.071 | -0.026 | 0.167 |
|  | PARIDERAS | 0.0017 | 0.0016 | 0.0018 | -0.161 | -0.249 | -0.072 |
|  | VILLALUENGA | 0.0017 | 0.0015 | 0.0018 | -0.043 | -0.127 | 0.041 |
|  | PAJARONCILLO | 0.0018 | 0.0017 | 0.0020 | -0.048 | -0.127 | 0.032 |
|  | SAN CLEMENTE | 0.0018 | 0.0017 | 0.0019 | -0.234 | -0.321 | -0.147 |
|  | PONFERRADA | 0.0021 | 0.0020 | 0.0023 | 0.086 | -0.002 | 0.174 |
|  | TORO | 0.0020 | 0.0018 | 0.0021 | -0.056 | -0.137 | 0.025 |
|  | MADRID | 0.0024 | 0.0022 | 0.0025 | -0.049 | -0.149 | 0.051 |
|  | RONDA | 0.0020 | 0.0018 | 0.0021 | 0.007 | -0.069 | 0.082 |
|  | CERRO DA CABEÇA | 0.0016 | 0.0015 | 0.0017 | 0.291 | 0.214 | 0.368 |
|  | CAZALLA | 0.0022 | 0.0020 | 0.0023 | 0.051 | -0.047 | 0.150 |
|  | GERENA | 0.0022 | 0.0021 | 0.0024 | 0.211 | 0.116 | 0.305 |
|  | GILENA | 0.0021 | 0.0020 | 0.0023 | -0.059 | -0.153 | 0.035 |
| **Mean** |  | **0.0020** | **0.0018** | **0.0021** | **0.005** | **-0.082** | **0.093** |
| Hercynian subgroup | CÓRDOBA | 0.0014 | 0.0013 | 0.0016 | 0.321 | 0.201 | 0.440 |
|  | SERRA DA ESTRELA | 0.0013 | 0.0012 | 0.0015 | 0.152 | 0.056 | 0.247 |
| **Mean** |  | **0.0014** | **0.0013** | **0.0016** | **0.236** | **0.128** | **0.343** |
| Algarve subgroup | FARO | 0.0017 | 0.0016 | 0.0019 | -0.07 | -0.178 | 0.038 |
|  | PINHEIROS DO MARIM | 0.0016 | 0.0015 | 0.0017 | 0.251 | 0.145 | 0.357 |
|  | QUARTEIRA | 0.0016 | 0.0014 | 0.0017 | -0.218 | -0.318 | -0.117 |
|  | VALE DO GARRAO | 0.0016 | 0.0015 | 0.0017 | -0.231 | -0.323 | -0.139 |
| **Mean** |  | **0.0016** | **0.0015** | **0.0018** | **-0.067** | **-0.168** | **0.034** |
| Doñana subgroup | PINAR ALGAIDA | 0.0016 | 0.0015 | 0.0017 | -0.183 | -0.291 | -0.074 |
|  | ABALARIO | 0.0016 | 0.0015 | 0.0018 | -0.313 | -0.423 | -0.203 |
|  | ALMONTE | 0.0017 | 0.0015 | 0.0018 | -0.560 | -0.667 | -0.453 |
|  | BONARES | 0.0016 | 0.0014 | 0.0017 | -0.502 | -0.613 | -0.390 |
|  | NORTH CARTAYA | 0.0015 | 0.0014 | 0.0016 | -0.136 | -0.232 | -0.041 |
|  | SOUTH CARTAYA | 0.0016 | 0.0015 | 0.0018 | -0.098 | -0.201 | 0.005 |
|  | HINOJOS | 0.0015 | 0.0014 | 0.0017 | -0.285 | -0.386 | -0.185 |
|  | NIEBLA | 0.0015 | 0.0013 | 0.0016 | -0.183 | -0.289 | -0.078 |
|  | EL PORTIL | 0.0018 | 0.0016 | 0.0019 | -0.032 | -0.125 | 0.061 |
|  | RIBETEHILOS | 0.0017 | 0.0015 | 0.0018 | -0.269 | -0.377 | -0.161 |
|  | VILLARRASA | 0.0018 | 0.0016 | 0.0019 | -0.077 | -0.169 | 0.015 |
|  | VILLAMANRIQUE | 0.0016 | 0.0015 | 0.0018 | -0.217 | -0.301 | -0.133 |
|  | LA JUNCOSILLA | 0.0016 | 0.0015 | 0.0017 | -0.368 | -0.475 | -0.261 |
| **Mean** |  | **0.0016** | **0.0015** | **0.0018** | **-0.247** | **-0.349** | **-0.146** |
| Cadiz  subgroup | LA BARROSA | 0.0016 | 0.0014 | 0.0017 | 0.025 | -0.083 | 0.132 |
|  | CHICLANA | 0.0015 | 0.0014 | 0.0017 | 0.256 | 0.164 | 0.349 |
|  | LAS LOMAS | 0.0013 | 0.0012 | 0.0014 | -0.005 | -0.105 | 0.094 |
|  | PUERTO REAL | 0.0016 | 0.0014 | 0.0017 | 0.136 | 0.025 | 0.247 |
|  | ROCHE | 0.0015 | 0.0014 | 0.0016 | -0.013 | -0.110 | 0.083 |
|  | EL SOTO | 0.0013 | 0.0012 | 0.0015 | 0.087 | -0.017 | 0.191 |
| **Mean** |  | **0.0015** | **0.0013** | **0.0016** | **0.081** | **-0.021** | **0.182** |
