## Supplementary material for "Patterns of Interploidy Admixture in Polyploid Complexes: Insights from *Thymus* sect. *Mastichina* (Lamiaceae)": Suppplemental Table S5

**Table S5**. Results of the ABBA-BABA test to quantify introgression events within *Thymus* sect. *Mastichina*.

| P1 | P2 | P3 | D | Z-score | p-value | f4-ratio | BBAA | ABBA | BABA |
| --- | --- | --- | --- | --- | --- | --- | --- | --- | --- |
| Algarve | **Hercynian** | **Tetraploid** | **0.061** | **7.237** | **<0.0001** | **1.00** | **131.9** | **115.2** | **101.8** |
| Doñana | **Hercynian** | **Tetraploid** | **0.046** | **5.047** | **<0.0001** | **1.00** | **134.2** | **112.5** | **102.5** |
| Cadiz | **Hercynian** | **Tetraploid** | **0.044** | **5.049** | **<0.0001** | **1.00** | **133.5** | **112.7** | **103.1** |
| Algarve | **Doñana** | **Tetraploid** | **0.015** | **2.761** | **0.006** | **0.37** | **132.2** | **109.9** | **106.6** |
| Algarve | **Cadiz** | **Tetraploid** | **0.018** | **2.349** | **0.019** | **0.42** | **130.4** | **110.8** | **107.0** |
| Doñana | Cadiz | Tetraploid | 0.002 | 0.298 | 0.766 | 0.07 | 136.1 | 106.8 | 106.3 |
| Doñana | **Hercynian** | **Algarve** | **0.020** | **2.674** | **0.007** | **0.28** | **115.2** | **110.2** | **105.8** |
| Cadiz | **Hercynian** | **Algarve** | **0.031** | **3.709** | **<0.001** | **0.37** | **115.4** | **111.3** | **104.7** |
| Doñana | Hercynian | Cadiz | 0.005 | 0.528 | 0.597 | 0.04 | 111.7 | 110.9 | 109.8 |
| Cadiz | Doñana | Algarve | 0.010 | 1.404 | 0.160 | 0.12 | 116.6 | 109.1 | 106.9 |

P1, P2, P3 refer to the three genetic groups used for the ABBA-BABA test, outgroup not shown
