## Supplementary material for "Patterns of Interploidy Admixture in Polyploid Complexes: Insights from *Thymus* sect. *Mastichina* (Lamiaceae)": Suppplemental Table S6

**Table S6**. Parameter estimates and model selection criteria are presented for four demographic scenarios (a-d) (see also Figure 7, and Table S5), each comparing diploids populations from the Algarve, Cádiz, Doñana and Hercynian regions, with the spatially closest population of the tetraploid *Thymus mastichina*.

| a) Tetraploid vs. Algarve |  |  |  |  | b) Tetraploid vs. Cádiz |  |  |  |
| --- | --- | --- | --- | --- | --- | --- | --- | --- |
| **Scenarios** | **AIC** | **deltaL** | **Ld** |  | **Scenarios** | **AIC** | **deltaL** | **Ld** |
| NO-GENE-FLOW | 7059.86 | 8.1359 | - |  | NO-GENE-FLOW | 1029.97 | 0.9209 | -309.6 |
| POP1_to_POP2 | 7062.31 | 8.2330 | - |  | POP1_to_POP2 | 1031.68 | 0.8580 | -309.5 |
| POP2_to_POP1 | 7030.06 | 1.2300 | -1524.45 |  | POP2_to_POP1 | 1031.74 | 0.8710 | -309.8 |
| BIDIRECTIONAL | 7031.85 | 1.1860 | **-1524.36** |  | BIDIRECTIONAL | 1031.80 | 0.4480 | **-308.8** |
| best_scenario | BIDIRECTIONAL | | |  | best_scenario | BIDIRECTIONAL | | |
| **Parameters** | **Mean** | **CI_Lower** | **CI_Upper** |  | **Parameters** | **Mean** | **CI_Lower** | **CI_Upper** |
| ANC_SIZE | 1114989 | 312095 | 1970655 |  | ANC_SIZE | 1178131 | 510340 | 1838977 |
| Split time | 37680 | 24961 | 48778 |  | Split time | 12305 | 8114 | 19113 |
| Ne1 (POP1) | 43565 | 33982 | 52339 |  | Ne1 (POP1) | 136518 | 80909 | 204574 |
| Ne2 (POP2) | 113893 | 42277 | 166116 |  | Ne2 (POP2) | 32964 | 25475 | 46187 |
| Mig Rate POP1→POP2 | 8.41e-05 | 6.37e-05 | 9.09e-05 |  | Mig Rate POP1→POP2 | 3.04e-05 | 1.07e-05 | 5.96e-05 |
| N_migr POP1→POP2 | 9586 | 7255 | 10353 |  | N_migr POP1→POP2 | 1002 | 1 | 1965 |
| Mig Rate POP2→POP1 | 2.59e-05 | 1.54e-07 | 8.99e-05 |  | Mig Rate POP2→POP1 | 6.89e-05 | 4.66e-05 | 8.89e-05 |
| N_migr POP2→POP1 | 1132 | 1 | 3916 |  | N_migr POP2→POP1 | 9 | 7 | 12 |
| MaxEstLhood | -15227 | -15314 | -15136 |  | MaxEstLhood | -19101 | -19212 | -18953 |
| MaxObsLhood | -15218 | -15303 | -15127 |  | MaxObsLhood | -18430 | -18540 | -18289 |
| c) Tetraploid vs. Doñana |  |  |  |  | d) Tetraploid vs. Hercynian | |  |  |
| **Scenarios** | **AIC** | **deltaL** | **Ld** |  | **Scenarios** | **AIC** | **deltaL** | **Ld** |
| NO-GENE-FLOW | 10949.31 | 44.5969 | - |  | NO-GENE-FLOW | 12091.08 | 50.8460 | - |
| POP1_to_POP2 | 10950.90 | 44.5079 | - |  | POP1_to_POP2 | 11891.31 | 7.0319 | **-2635.3** |
| POP2_to_POP1 | 10826.65 | 17.5269 | -2368.5 |  | POP2_to_POP1 | 11978.04 | 25.8640 | - |
| BIDIRECTIONAL | 10800.25 | 11.3589 | **-2362.9** |  | BIDIRECTIONAL | 11895.52 | 7.5119 | -2635.6 |
| best_scenario | BIDIRECTIONAL | | |  | best_scenario | POP1 to POP2 | | |
| **Parameters** | **Mean** | **CI_Lower** | **CI_Upper** |  | **Parameters** | **Mean** | **CI_Lower** | **CI_Upper** |
| ANC_SIZE | 2421642 | 1722269 | 3138319 |  | ANC_SIZE | 2663848 | 2073080 | 3847639 |
| Split time | 44805 | 39374 | 50352 |  | Split time | 36616 | 32362 | 40296 |
| Ne1 (POP1) | 111194 | 95003 | 125720 |  | Ne1 (POP1) | 64376 | 58299 | 71467 |
| Ne2 (POP2) | 54570 | 41643 | 67274 |  | Ne2 (POP2) | 39769 | 36676 | 42523 |
| Mig Rate POP1→POP2 | 8.75e-05 | 8.35e-05 | 9.09e-05 |  | Mig Rate POP1→POP2 | 8.69e-05 | 8.18e-05 | 9.08e-05 |
| N_migr POP1→POP2 | 4775 | 4557 | 4959 |  | N_migr POP1→POP2 | 3456 | 3253 | 3611 |
| Mig Rate POP2→POP1 | 3.09e-05 | 1.38e-05 | 4.89e-05 |  | Mig Rate POP2→POP1 | 5.90e-07 | 1.67e-07 | 1.46e-06 |
| N_migr POP2→POP1 | 3436 | 1534 | 5437 |  | N_migr POP2→POP1 | 0.0 | 0.0 | 1 |
| MaxEstLhood | -23617 | -23744 | -23507 |  | MaxEstLhood | -26410 | -26538 | -26302 |
| MaxObsLhood | -23507 | -23633 | -23400 |  | MaxObsLhood | -26337 | -26450 | -26223 |
| Ld=Likelihood distributions; CI=Confidence Interval | | |  |  |  |  |  |  |
