## Supplementary figures and images for "Patterns of Interploidy Admixture in Polyploid Complexes: Insights from *Thymus* sect. *Mastichina* (Lamiaceae)"

### Suppplemental Figure S1

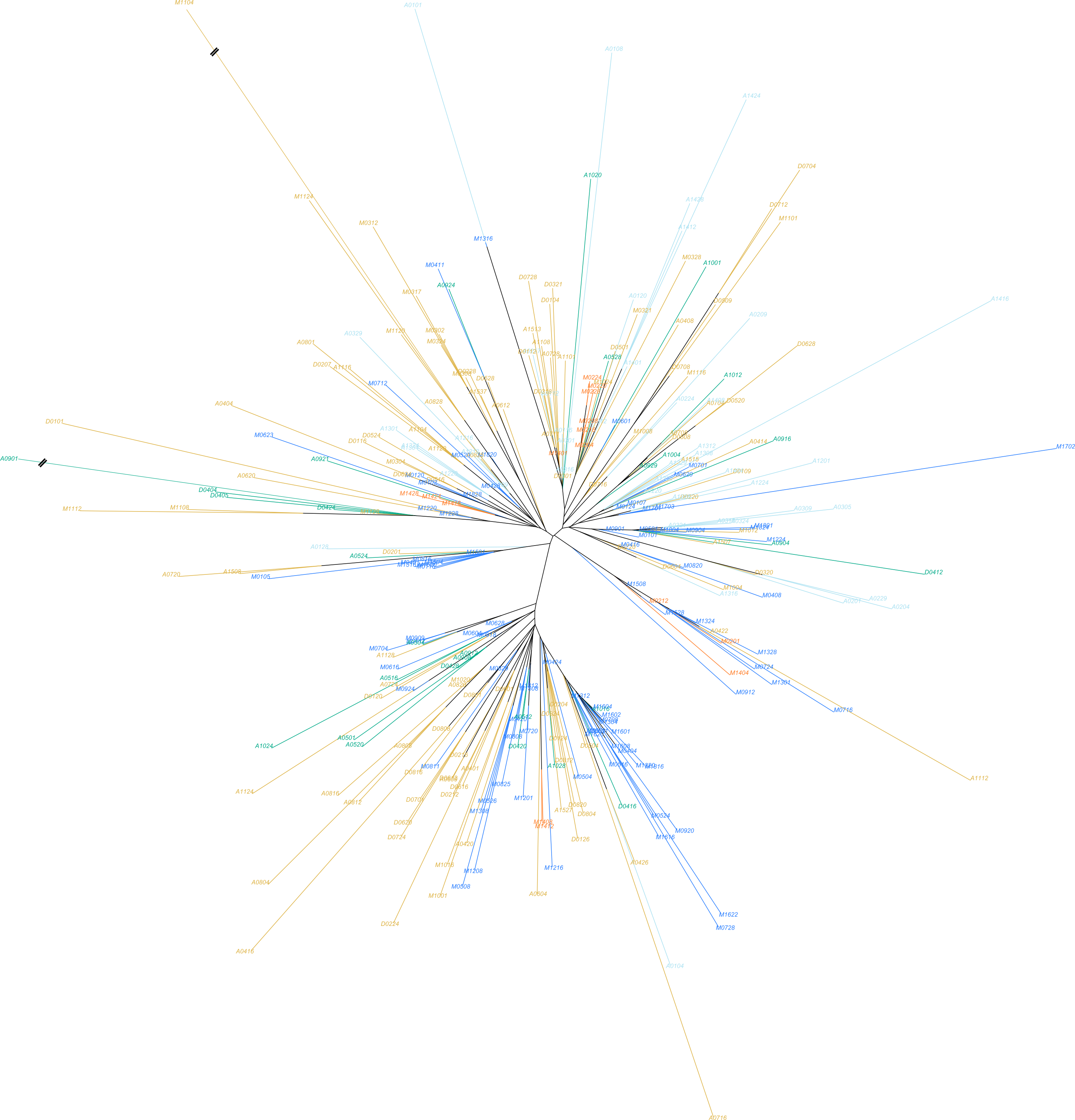

### Suppplemental Figure S2

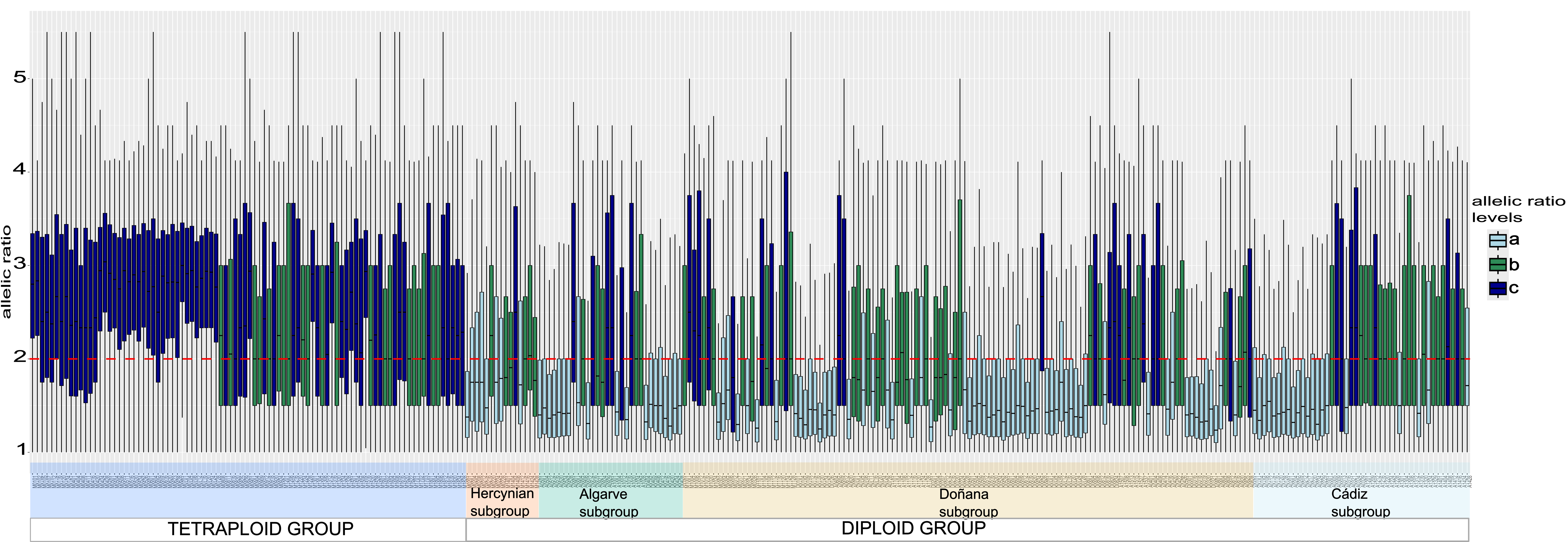

### Suppplemental Figure S3

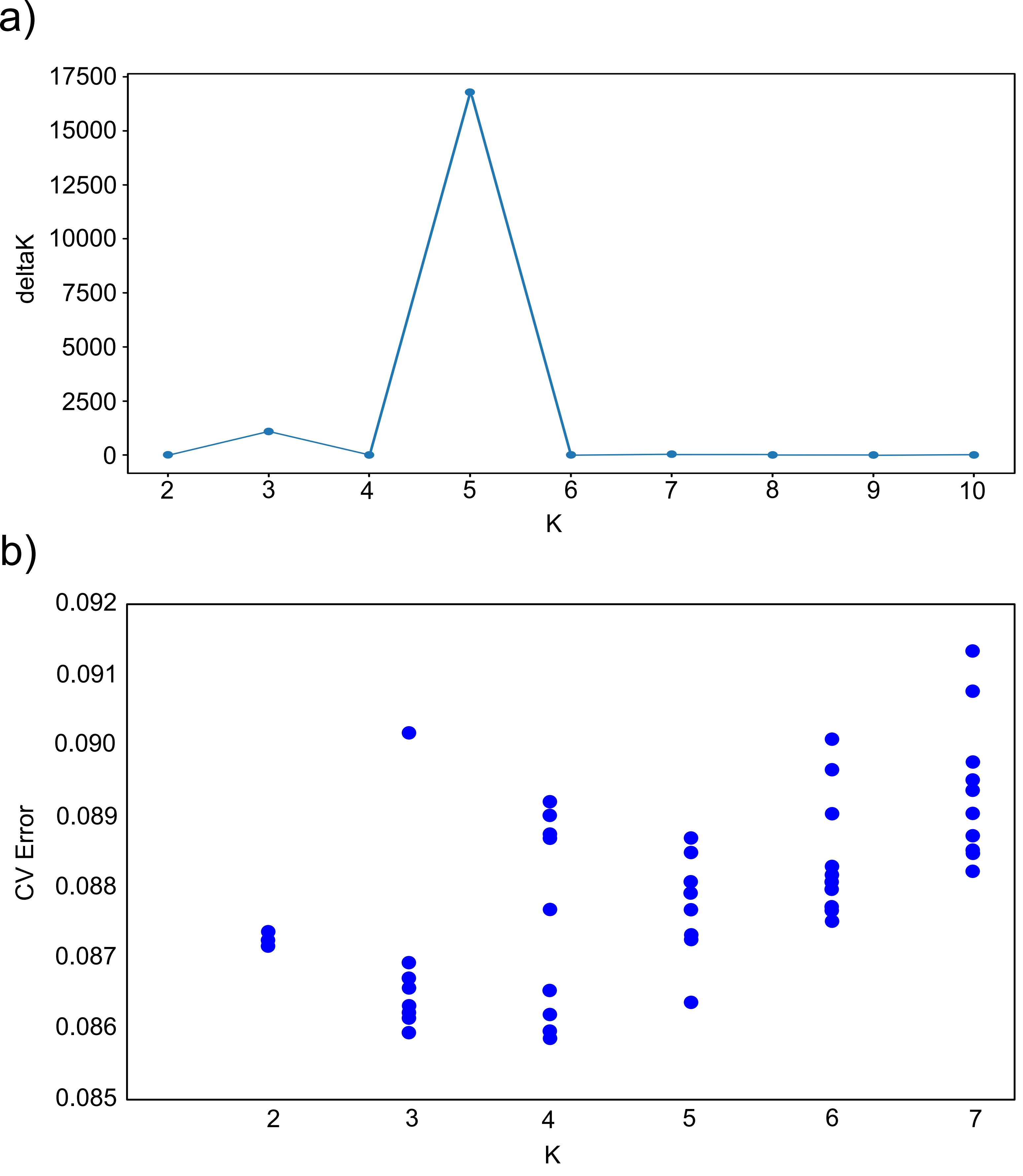

### Suppplemental Figure S4

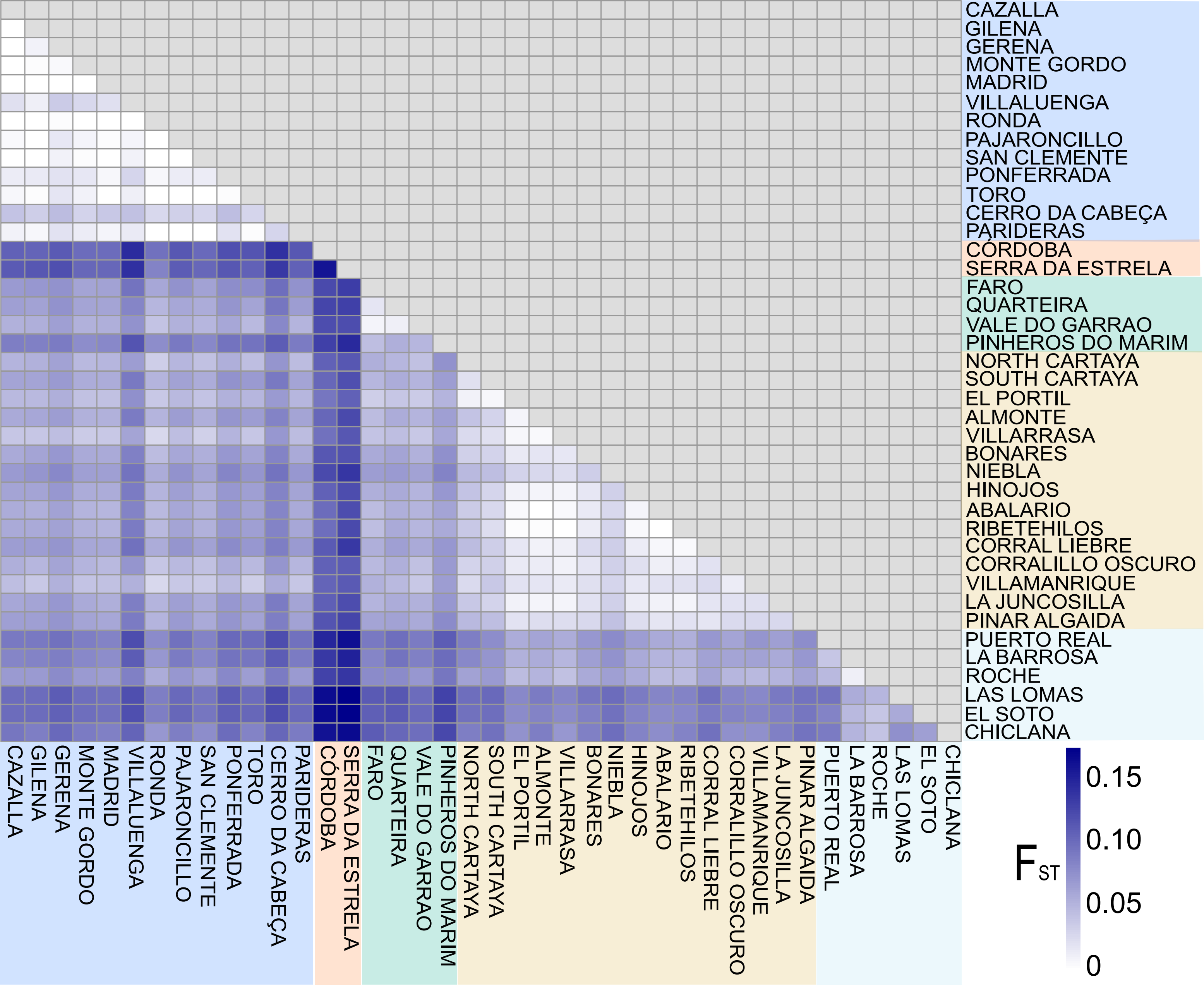

### Suppplemental Figure S5

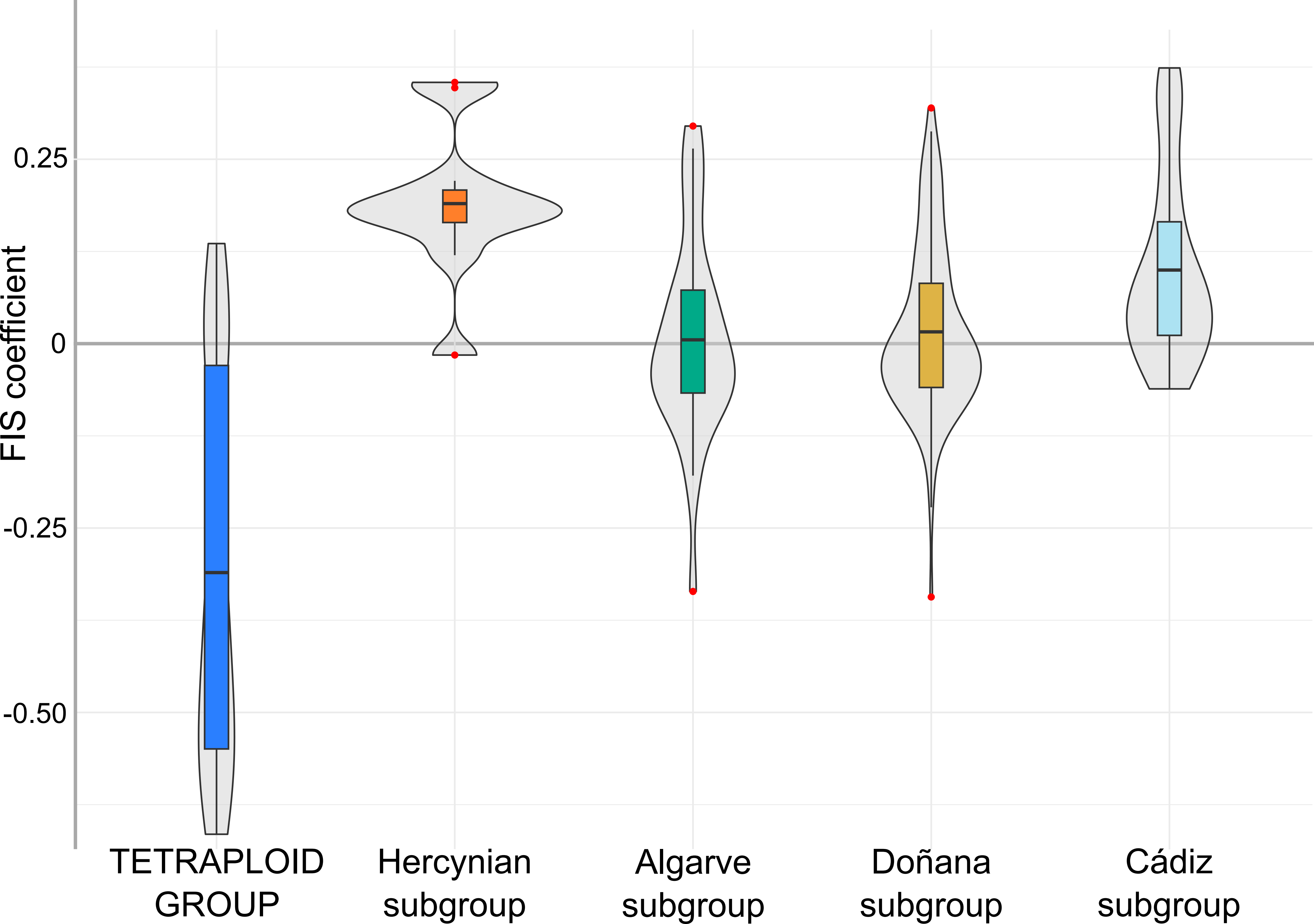

### Suppplemental Figure S6

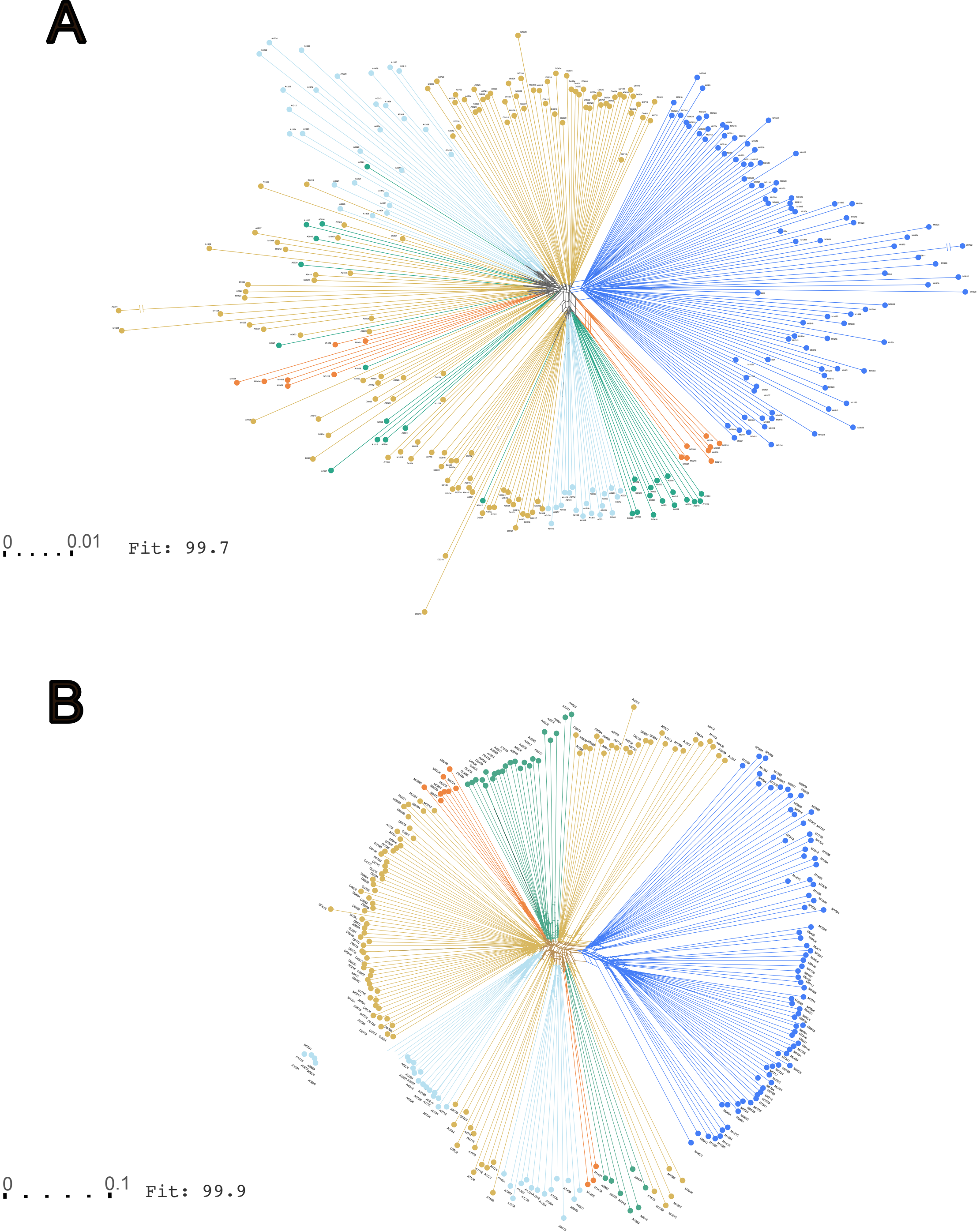
